# Benchmarking sequence-to-function models and mapping enhancer–gene interactions with E2GScore

**DOI:** 10.64898/2026.11.24.769033

**Authors:** Caleb The-Tjoean, Hamid Alinejad-Rokny, Irina Voineagu

## Abstract

Sequence-to-function models predict gene expression and chromatin features directly from DNA sequence and are increasingly used to infer enhancer–gene regulatory interactions. However, differences in attribution procedures and benchmarking datasets complicate comparisons between models, while the biological information captured by their enhancer–gene scores remains poorly understood. Here we introduce E2GScore, a unified framework for extracting and benchmarking enhancer–gene predictions from sequence-to-function models. By systematically varying scoring protocols, we identify gene-readout normalization and cell-type-matched expression tracks as important determinants of performance, whereas in silico mutagenesis offers no advantage over less computationally demanding input gradients. We compare seven sequence-to-function models and three established enhancer–gene predictors against CRISPR interference screens in K562 cells and primary human astrocytes. AlphaGenome 1 and 2 outperformed all other sequence-to-function models, and matched or outperformed experimentally informed predictors, particularly for distal enhancers, without a statistically significant difference between AlphaGenome versions after multiple-testing correction. We further show that the apparent reduction in predictive performance in astrocytes relative to K562 cells observed across models results primarily from differences in benchmarking dataset composition, particularly enhancer–TSS distance and target-gene expression. Using the open-weight AlphaGenome 1, we then generated E2GMap, a genome-wide resource of enhancer–gene regulation comprising 14.7 billion enhancer–gene attribution scores across 454 human biosamples, 45% of which are absent from existing enhancer–gene maps such as those from ENCODE-rE2G and AlphaGenome 2. E2GMap is available at e2gscore.com. Together, E2GScore and E2GMap establish a framework for rigorous evaluation of sequence-based enhancer–gene predictions and provide an extensible resource for studying gene regulation across human cell types and tissues.

## Introduction

Most genetic variants associated with human disease lie in non-coding DNA and are concentrated in regulatory elements such as enhancers [1,2]. Interpreting these variants requires knowing which genes each enhancer regulates, but enhancers can act over hundreds of kilobases and frequently do not regulate the nearest gene [3,4]. CRISPR interference (CRISPRi) screens, which silence candidate enhancers and measure the effect on gene expression, provide direct evidence of enhancer–gene regulation [5–11]. However, such screens test only thousands of enhancer–gene pairs and have been carried out at scale in few cell types. Computational prediction is therefore needed to extend enhancer–gene maps across the genome and across cell types. The leading predictors, including the activity-by-contact (ABC) model [5,8], ENCODE-rE2G [9], and EGrf [10], combine chromatin accessibility of the cell type of interest with histone modification or chromatin contact data, and the latter two are trained directly on CRISPRi data.

Sequence-to-function (seq2func) models, by contrast, are deep neural networks trained to predict genome-wide functional genomics measurements, such as chromatin accessibility, histone modifications, and gene expression, directly from DNA sequence across hundreds of cell types and tissues [12–18].

Seq2func models do not, by design, predict enhancer–gene regulation. Given a DNA sequence they predict signal tracks. Predicting whether an enhancer regulates a gene requires an additional attribution procedure [19] that estimates how much the enhancer’s sequence contributes to the gene’s predicted expression signal. Three sets of decisions shape the resulting enhancer–gene linking prediction. **(i)** *Attribution method*. Input gradients measure how sensitive the gene’s predicted expression is to each nucleotide of the input sequence, and the enhancer’s score summarizes these values across the element [20]; in silico mutagenesis (ISM) instead perturbs the sequence of the whole enhancer and measures the resulting change in predicted expression [21]. **(ii)** *Gene readout*. Either cap analysis of gene expression (CAGE) or RNA-seq can be used as the assay representing the gene’s expression signal. In addition, the expression signal can be summarized at a single transcription start site (TSS), at all annotated TSSs of a gene, or across the gene’s exons. **(iii)** *Input sequence*. The model’s window can be centered on the TSS or on the gene midpoint. Successive models have widened the sequence context they consider, from 197 kb in Enformer [22] to 524 kb in Borzoi [23] and 1 Mb in AlphaGenome [24,25]. Thus the choice of how to center the input sequence, together with each model’s fixed window length, determines which enhancers can be scored for regulatory effects.

In the published evaluations these decisions are made differently, so their reported accuracies cannot be compared directly, and it is unclear how much of the difference between models reflects the models themselves rather than the scoring procedure [26,27]. Independent assessments have also shown that conclusions about seq2func models depend strongly on how the models are evaluated. Enformer captures promoter-proximal determinants of expression but underestimates the effects of distal enhancers measured in CRISPRi screens [28], and models that accurately predict differences in expression between genes explain little of the variation in expression between individuals [29,30]. Murphy and Koo [31] recently showed that seq2func models systematically underpredict the effect size of enhancer perturbation on gene expression.

Furthermore, enhancer–gene prediction by seq2func models has been evaluated primarily on CRISPRi data from K562 [32], which is also among the cell types best represented in the models’ training data. Whether performance estimated in K562 transfers to primary human cells has therefore been unclear. We recently reported a CRISPRi screen of candidate enhancers in primary human astrocytes [10], which provides a benchmarking dataset of well-powered pairs outside immortalized cell lines. Two recent preprints have used this screen to evaluate AlphaGenome: Huang et al. [33] found that AlphaGenome enhancer-deletion responses add predictive value to a supervised predictor, and Gayoso et al. [25] reported that AlphaGenome 2 outperforms supervised predictors in astrocytes. In the latter study, however, accuracy, measured as the area under the precision–recall curve (auPRC), was lower on the astrocyte screen (auPRC 0.68) than on K562 cells (auPRC ≈0.80; their Supplementary Fig. 11f), and whether such differences reflect weaker generalization across cell types or differences between the benchmarking datasets themselves has not been examined. Earlier work on predicting enhancer–promoter interactions likewise showed that the design of a benchmark can inflate the apparent performance of the first predictors [34–38].

Here we present E2GScore, a framework that makes every step of enhancer–gene scoring an explicit parameter [39], so that models can be compared under matched protocols. After confirming that E2GScore reproduces the published benchmarks [22–24], and those of AlphaGenome 2 through its API [25], we evaluate 10 models on the K562 and astrocyte CRISPRi screens. These include seq2func models: Enformer [22], Borzoi [23], Flashzoi [40], Nucleotide Transformer v3 (NTv3) [41], Cerberus [42], AlphaGenome 1 [24], and AlphaGenome 2 [25]; and expert models: ABC [5], ENCODE-rE2G (DNase and extended versions) [9], and EGrf (standard and extended versions) [10].

We used confidence intervals (CIs) from bootstrapping over groups of pairs that share an enhancer or a gene, which accounts for their dependence, and paired tests between methods on the same pairs.

We first show that normalization across genes, cell-type context, and the TSS used for CAGE matter, and that ISM gives no advantage over input gradients. We further show that model performance is generally robust to the assay used as readout (CAGE or RNA-seq), and that a substantial part of the lower performance in astrocytes is attributable to the composition of the benchmarking dataset. We also show, with paired statistical comparisons, that AlphaGenome 1 and 2 match the best supervised predictors in K562 and match or outperform them in astrocytes, in line with the astrocyte results recently reported for AlphaGenome 2 [25]. In our evaluation the performance of AlphaGenome 2 did not differ significantly from that of AlphaGenome 1 for enhancer–gene linking on either screen.

We further present E2GMap, a collection of genome-wide enhancer–gene maps employing AlphaGenome 1’s attributions across 454 biosamples, complementing the ENCODE-rE2G [9] and AlphaGenome 2 [25] maps.

## Results

### 1. E2GScore, a unified enhancer–gene scoring framework

We developed E2GScore, a unified framework for scoring and evaluating seq2func models on enhancer– gene interaction prediction (Fig. 1a). E2GScore makes 42 methodological choices explicit and configurable, enabling systematic evaluation of enhancer–gene attribution scores derived from seq2func models using input gradients or whole-element ISM (Methods; Supplementary Table 1). We call a given configuration of these choices a “protocol”. To verify that E2GScore correctly implements each model and its published protocol, we first reproduced published enhancer–gene linking benchmarking results for Enformer [22] (Fulco and Gasperini screens [5,6]), Borzoi [23] (Gasperini and Nasser screens [6,8]), and AlphaGenome 1 [24] (Gschwind screen [9]) (Supplementary Figs. 1–4), across a range of protocols. Our auPRCs (computed as average precision, with 95% CIs from bootstrapping pairs in each distance bin; Methods) matched the published values for Enformer (38 of 40 within our 95% CI; median absolute difference 0.0124), Borzoi (24 of 24; median absolute difference 0.0004), and AlphaGenome 1 (5 of 5; median absolute difference 0.0055). The benchmarks of AlphaGenome 2 [25] were reproduced through its API (24 of 24; median absolute difference 0.0028; Supplementary Fig. 4c–e).

**Fig. 1.**
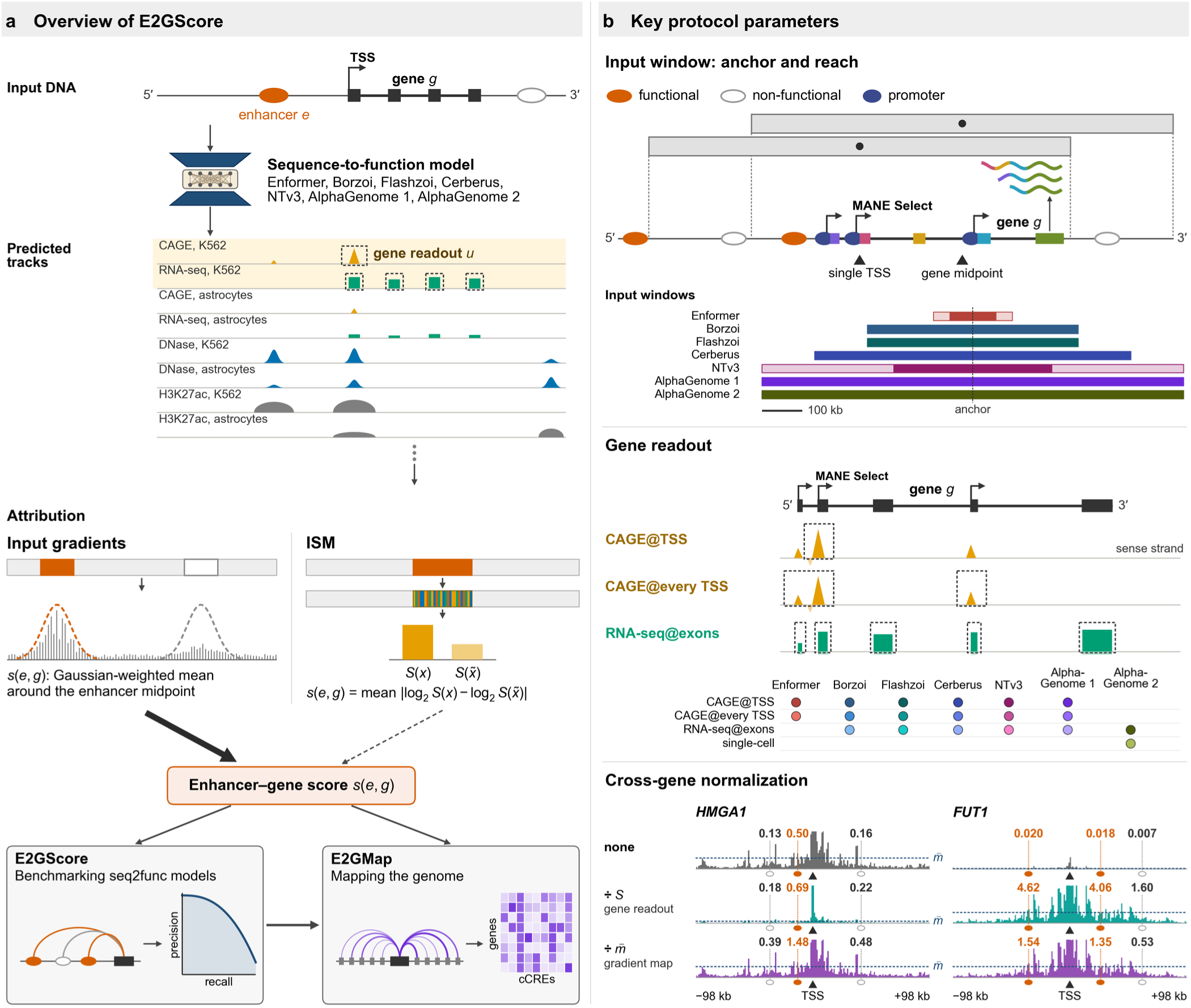
E2GScore makes each step of enhancer–gene scoring an explicit choice. **a**, A sequence-to-function (seq2func) model predicts tracks for one input window; the gene readout *u* (**b**) is attributed to a candidate enhancer by input gradients (Gaussian-weighted mean of |gradient × input| at the enhancer; σ = 300 bp, cut at 4σ) or by in silico mutagenesis (ISM; mean absolute change in log_2_ *S* between the reference sequence *x* and each of *N* shuffles *x̃* of the enhancer). Arrows: thick, used for E2GScore and E2GMap (Figs. 2–4); dashed, used for E2GScore but not for E2GMap (Supplementary Fig. 10). Input gradients score all enhancers of a gene in one forward and backward pass, whereas ISM needs *N* forward passes per enhancer–gene pair and one reference pass per gene; scores are evaluated against CRISPRi labels (E2GScore; Figs. 2 and 3) and, with AlphaGenome 1, computed genome-wide for protein-coding genes and ENCODE4 candidate cis-regulatory elements (cCREs; E2GMap; Fig. 4). **b**, Key protocol parameters. Input window: centered on a single TSS (e.g. MANE Select) or the gene midpoint; elements outside it score 0. CAGE@TSS reads the TSS on which its window is centered; CAGE@every TSS and RNA-seq@exons use the gene midpoint. Bars, input windows to scale (197 kb, Enformer; 524 kb, Borzoi and Flashzoi; 786 kb, Cerberus; 1 Mb, NTv3, AlphaGenome 1, and AlphaGenome 2); pale, input outside the predicted region (Enformer and NTv3 predict only the central 114 kb and 393 kb, respectively). Gene readout: *u* = log *S*, *S* = Σ_b∈B_ ȳ_b_, where *B* is the set of output bins of the region (384 bp at the TSS or at every TSS, merged where they overlap, or exon bins) and ȳ_b_ the mean over the cell type’s tracks; stranded tracks are read on the gene’s sense strand. Dots, the 19 readouts of Fig. 2 in its colors (single-cell, AlphaGenome 2’s single-cell expression head). Cross-gene normalization: Enformer maps of |gradient × input| of the summed readout for two K562 genes (CAGE at the K562 table’s RefSeq TSS; 1-kb means), raw (none) or divided by the gene readout (*S*; scores ×10^3^) or the mean absolute input gradient (m̅); numbers, scores of marked elements (orange, functional; white, non-functional).

We next used E2GScore to vary key protocol parameters while holding all others fixed, and evaluated the effects on enhancer–gene linking prediction using Enformer, Borzoi, and AlphaGenome 1 in K562 and astrocytes (NHA) (Fig. 1b) with ground-truth data from Gschwind et al. [9] and Green et al. [10]. The K562 and astrocyte screens include 471/10,356 and 133/2,307 positive/total enhancer–gene pairs, respectively.

Several parameter choices significantly changed model performance (Supplementary Figs. 5–8), in particular cross-gene normalization, cell-type context for the expression readout, and the TSS for the CAGE protocol.

Without normalization across genes, the enhancers of highly expressed genes simply outranked those of weakly expressed genes. Supplementary Figs. 5a and 6a show examples of how the lack of normalization across genes significantly worsens performance for both input gradients and ISM. For input gradients, genereadout normalization was better than no normalization and in most cases better than gradient-map normalization, while using both was equivalent to gradient-map normalization alone (empirically and by identity): the cancellation was exact for Enformer and AlphaGenome under all protocols (auPRC within 2 × 10⁻⁵) and approximate for Borzoi, since each of its replicates’ maps was divided by its own mean absolute input gradient before averaging (Supplementary Fig. 5b,c). Likewise, for ISM, cross-gene normalization improved predictions across all models in all paired comparisons (Supplementary Fig. 6b,c).

Using gene expression readouts from tracks of the cell type studied, rather than from tracks of all or random cell types, improved all three models in K562; without cell-type-matched tracks Enformer and Borzoi performed barely better than enhancer–TSS distance (Supplementary Fig. 7).

For the CAGE readout we assessed whether the TSS annotation mattered. Because the CAGE signal is read at the chosen TSS and the input window is centered on it, each choice changes the gene readout and the window anchor at once. We compared the TSS from the Matched Annotation from NCBI and EMBL-EBI (MANE) Select transcript [43], the 5′-most annotated TSS, the RefSeq TSS shared by the most coding transcripts of the gene, the major TSS using the predicted CAGE tracks, and the major TSS using the measured CAGE signal (Supplementary Fig. 8a–i). We found the MANE Select TSS and the major predicted/measured TSS were the best TSSs in both screens, on par with CAGE signal aggregated over all annotated TSSs, except for Enformer in K562 (Fig. 2c,d; Supplementary Fig. 8j).

**Fig. 2.**
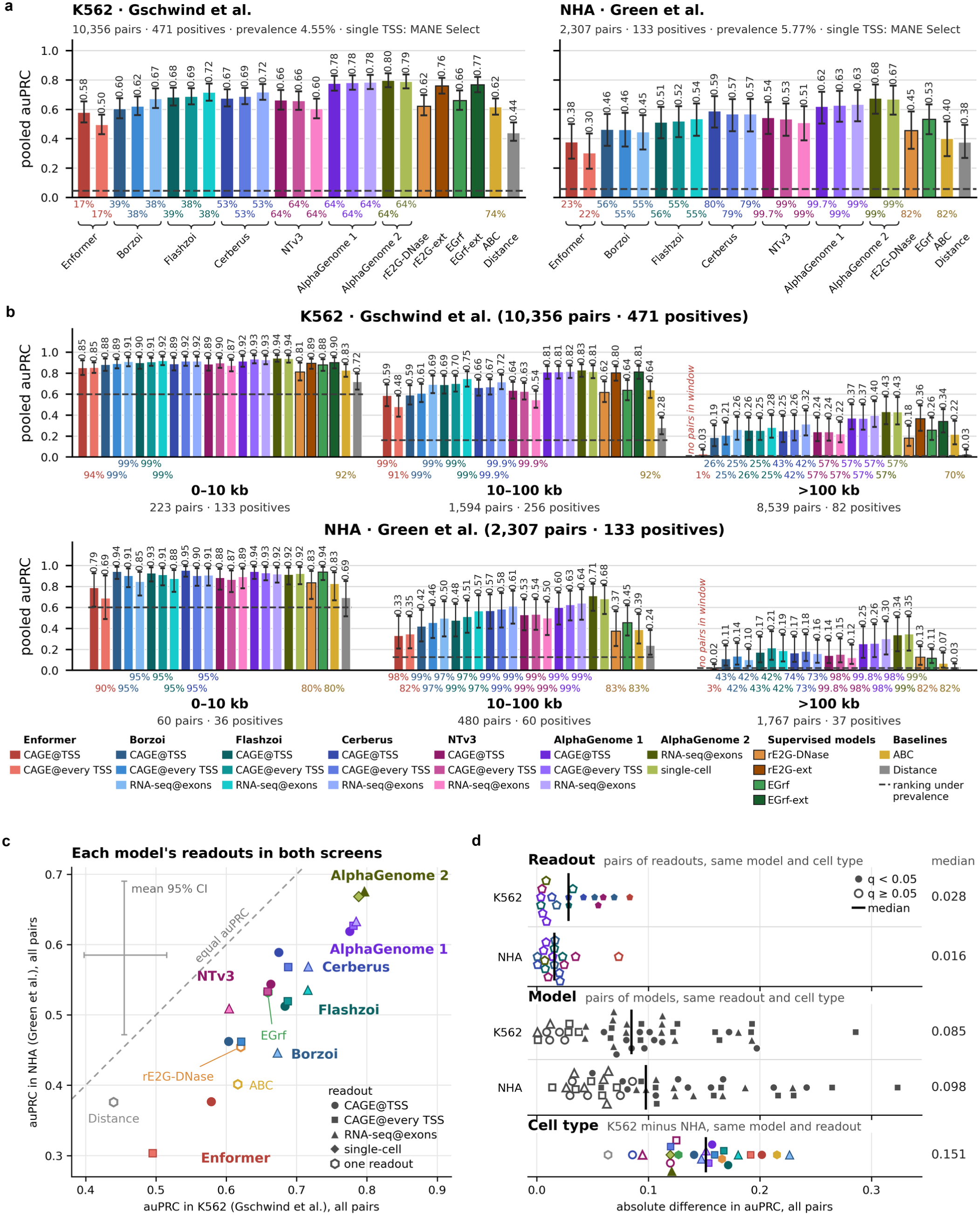
Under matched protocols, model rankings for gradient-based attribution scores are consistent across gene readouts and cell types. **a,** Pooled auPRC over all pairs of the K562 [9] (10,356 pairs, 471 positives) and astrocyte (NHA; [10]; 2,307 pairs, 133 positives) screens. Seq2func models (color, model; shade, readout) are scored by input gradients, AlphaGenome 2 through its API; ENCODE-rE2G and EGrf are out-of-fold predictions for held-out chromosomes (extended versions, K562 only); ABC, activity-by-contact; Distance, 1/distance to the MANE Select TSS. Percentages, share of pairs a method can score (others take 0); dashed lines, random ranking under prevalence. **b,** As a, by distance to the MANE Select TSS. Error bars, 95% confidence interval (CI; 10,000 bootstrap resamples of connected components). **c,** auPRC in NHA against K562; shape, readout; hexagons, single-readout methods; cross, mean 95% CI. **d,** Absolute auPRC difference over all pairs between a model’s readouts (*n* = 17 per screen), models with the same readout (*n* = 45 per screen), and screens (*n* = 23). Filled, Benjamini–Hochberg *q* < 0.05; tick, median.

Other parameter choices, including the input window anchor, shape of the Gaussian kernel, Gaussian or average pooling, and gradient correction, had minor or non-significant effects on performance (Supplementary Fig. 9). Narrowing the CAGE readout from 384 bp to 128 bp lowered auPRC for Enformer CAGE@TSS and Borzoi (significant in three of their six comparisons, all in K562) but not for AlphaGenome 1 (Supplementary Fig. 9). Moreover, perturbation-based attribution scores (ISM) performed worse than gradient-based scores (input gradients) in K562, did not differ significantly from them in astrocytes, and also required substantially more compute, linearly scaling with the number of enhancer–gene pairs rather than the number of genes (Supplementary Fig. 10). Caching the convolutional activations sped up ISM by 2.1–5.2× in test runs without changing any pair’s rank, but its cost still scaled with the number of pairs (Methods).

We therefore fixed the following parameters for subsequent model comparisons: input gradients with genereadout normalization, using tracks from the cell type studied, and the MANE Select TSS for CAGE (Methods; Supplementary Table 1).

### 2. Under matched protocols, model rankings are consistent across gene readouts and cell types

Next we asked how well the models perform across cell types, by comparing enhancer–gene predictions in K562 cells with predictions in astrocytes, a cell type that is far less represented in the models’ training data than K562. We used E2GScore to benchmark seven seq2func models (Enformer, Borzoi, Flashzoi, NTv3, Cerberus, AlphaGenome 1, and AlphaGenome 2) (Methods), three expert models (ABC, ENCODE-rE2G, and EGrf) (Methods), and enhancer–TSS distance as a baseline (Fig. 2).

For each model we compared up to three gene readouts: CAGE signal at a single representative TSS (that of the MANE Select transcript; CAGE@TSS), CAGE signal summed over all of the gene’s annotated TSSs (CAGE@every TSS), and RNA-seq coverage summed over the gene’s exons (RNA-seq@exons). AlphaGenome 2 was read from RNA-seq@exons and, as an additional readout, from its single-cell expression head. Model performance was measured as the auPRC over all tested pairs of a screen. Differences between protocols or models are reported as the paired difference in auPRC on the same pairs, with its 95% CI and Benjamini–Hochberg q value (Methods; [44]; all comparisons are listed in Supplementary Table 2).

The ranking of seq2func models by auPRC over all pairs was consistent across readouts and in both screens: overall AlphaGenome 2 and AlphaGenome 1 performed best, followed by Cerberus, Flashzoi, and NTv3, then Borzoi and Enformer (Fig. 2a,c). These differences arose almost entirely from distal enhancers. Within 10 kb of the TSS all models performed similarly, whereas beyond 10 kb they separated widely (Fig. 2b).

The differences between two seq2func models for distal enhancers could reflect how well each model ranked the pairs both could score, or the size of their input window. To separate these, we split each model’s auPRC gap to AlphaGenome 1 into two parts: ranking, the difference in auPRC on the pairs both models can score, and reach, the remainder of the auPRC gap. Across the different readouts and CRISPRi screens, reach accounted for 1–56% of AlphaGenome 1’s advantage over Cerberus, 8–51% over Flashzoi, 5–27% over Borzoi, and 27–66% over Enformer (whose 197-kb window holds only 60–83% of positives; Supplementary Figs. 11 and 12). NTv3 and AlphaGenome 1 can score almost exactly the same pairs (all but one K562 pair under CAGE@every TSS), so the gap between them is almost entirely ranking; models that had lower reach generally had higher auPRC penalties (Supplementary Fig. 11). These data show that most of AlphaGenome 1’s advantage over other methods reflected better ranking of the pairs both methods could score, rather than the wider input window.

When each gene’s enhancers, or each enhancer’s genes, were ranked separately, no single method led, likely due to limited sample size in these comparisons (Supplementary Fig. 13b,c).

The choice of gene readout had little effect on performance (Fig. 2c,d; Supplementary Fig. 10c,d). Readouts of the same model and cell type differed by a median of 0.028/0.016 auPRC in K562/astrocytes, compared with 0.085/0.098 between models under the same readout and cell type, and 0.151 between the two cell types for the same model and readout. No readout comparison was significant in astrocytes (Fig. 2d; Supplementary Fig. 10c,d). In K562, RNA-seq outperformed CAGE for Borzoi, Cerberus, and Flashzoi, but was worse than CAGE for NTv3; Enformer performed worse with CAGE@every TSS than CAGE@TSS since centering its 197-kb input on the gene midpoint reaches fewer enhancers.

When seq2func models were compared to expert models, AlphaGenome performed as well as or better than the expert models. In K562, AlphaGenome 1 and 2 matched or exceeded the extended versions of ENCODE-rE2G and EGrf (which utilize K562 assays not available for other cell types) (Fig. 2a). In astrocytes, both AlphaGenome versions, with the RNA-seq readout, outperformed every expert model, including the best, EGrf, with the advantage coming from enhancers more than 10 kb from the TSS (Fig. 2b), in agreement with Gayoso et al. [25]. This advantage was specific to AlphaGenome: none of the other seq2func models significantly outperformed EGrf in astrocytes.

We also compared the two versions of AlphaGenome. In K562, AlphaGenome 2 led AlphaGenome 1 by 0.012 auPRC (P = 0.28; q = 0.36) using the RNA-seq readout for both models, and by 0.004 auPRC (P = 0.76; q = 0.81) using the single-cell RNA-seq (scRNA-seq) readout for AlphaGenome 2 and RNA-seq for AlphaGenome 1. In astrocytes, AlphaGenome 2 led AlphaGenome 1 by 0.043 auPRC (P = 0.039; q = 0.080) using the RNA-seq readout for both models, and by 0.035 auPRC (P = 0.071; q = 0.13) using the scRNA-seq readout for AlphaGenome 2 and RNA-seq for AlphaGenome 1. These data showed that AlphaGenome 1 with its best readout (RNA-seq) did not differ significantly from AlphaGenome 2 on either screen, after correction for multiple comparisons, although the point estimate was higher for the latter version. Nominally AlphaGenome 2 outperformed AlphaGenome 1 in astrocytes only when both models used the RNA-seq readout; we note that AlphaGenome 2’s astrocyte condition was trained on single-cell data from the screen’s own cells [25].

### 3. Benchmarking dataset composition confounds apparent model performance primarily via enhancer–TSS distance

Every method, including the expert models, performed worse on the astrocyte screen than on K562, by a median of 0.151 auPRC (Fig. 2c,d; Supplementary Table 2). This could reflect poorer prediction of the gene expression readout in astrocytes, poorer generalization of enhancer–gene linking to a primary cell type, or differences between the two benchmarking datasets. We confirmed that the seq2func models (Enformer, Borzoi, and AlphaGenome 1) predicted gene expression at least as accurately in astrocytes as in K562 (Supplementary Fig. 14).

We therefore asked whether the composition of the two benchmarking datasets accounts for the performance gap. The screens differ in three properties that affect auPRC independently of model quality: the number and fraction of tested pairs that are positive (prevalence), the distribution of enhancer–TSS distances, and the expression of target genes (Fig. 3a,b). The astrocyte screen strictly excluded elements less than 2 kb from a gene, and tested effects within 500 kb, while the K562 screen included proximal elements and tested effects up to 2.4 Mb (distances to each screen’s own TSS; Fig. 3a). The distributions of enhancer–TSS distances between the K562 and astrocyte screens differed more than the distributions of target-gene expression levels between screens did (Kolmogorov–Smirnov D = 0.37, q < 0.001, against D = 0.18, q = 0.023); this was mainly the case for negative pairs in both screens (Fig. 3a,b), reflecting differences in the design of each screen’s tested pairs.

**Fig. 3.**
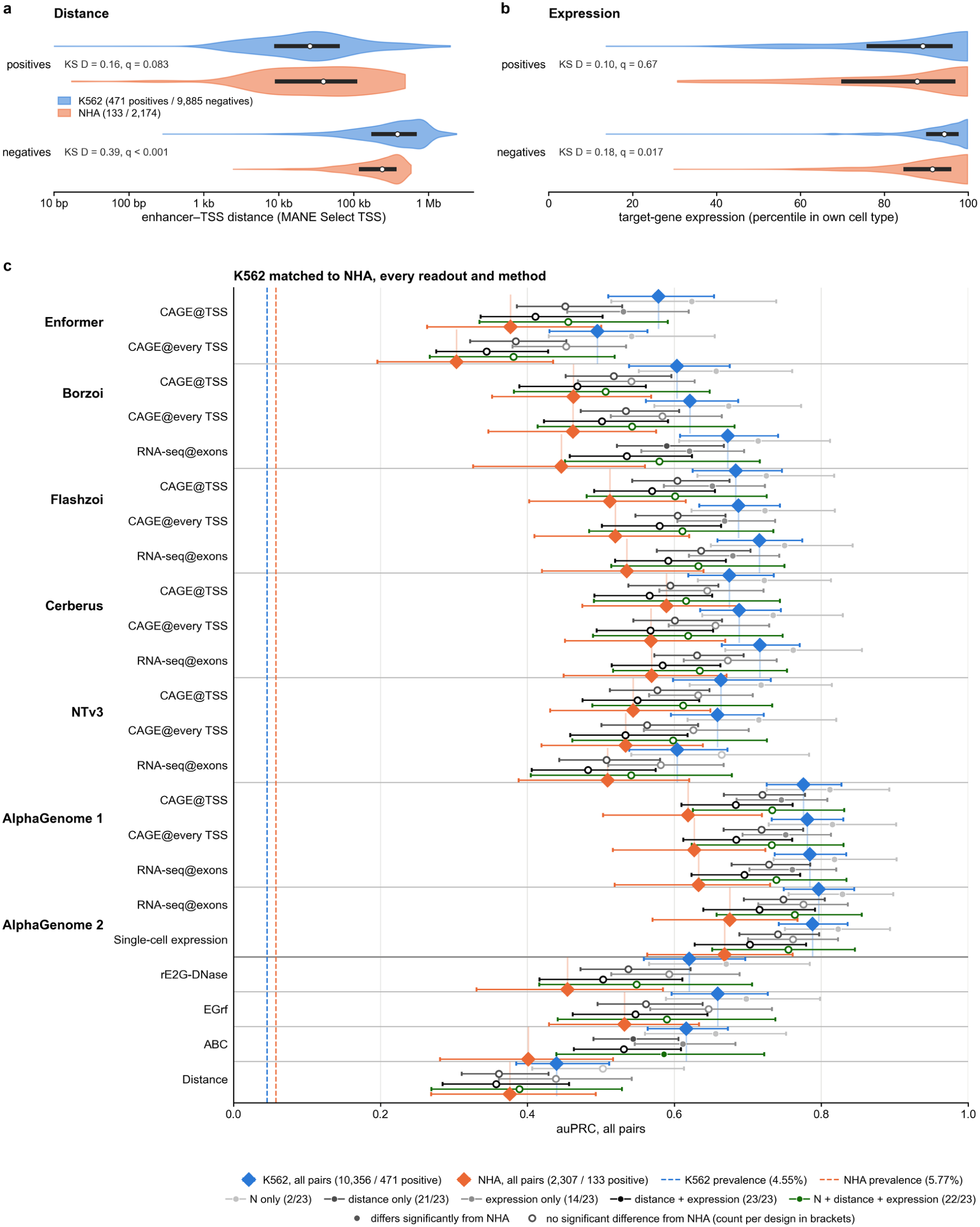
Benchmark composition, primarily enhancer–TSS distance, confounds the comparison of auPRC between K562 and astrocytes. **a**,**b**, Enhancer–TSS distance (**a**; to the MANE Select TSS) and target-gene expression (**b**; percentile within the gene’s own cell type) of the positive and negative pairs of K562 (blue; 471 and 9,885) and astrocytes (NHA, orange; 133 and 2,174); in **b**, 9,879 (K562) and 2,170 (NHA) negatives have an expression percentile. Violins, kernel densities; white dot, median; black bar, interquartile range; KS D, two-sample Kolmogorov–Smirnov distance between the screens; q, Benjamini–Hochberg-adjusted P value over six comparisons (these four and the two over all pairs), from 10,000 permutations of the screen label over connected components of genes and elements (Supplementary Table 4). **c**, auPRC over all pairs for 19 readouts of seven seq2func models, three expert models, and distance: K562 (blue diamonds), NHA (orange diamonds), and K562 matched to NHA (circles) in the numbers of positive and negative pairs (N; mean of 10 subsamples), distance, expression, or combinations of these. Open circles, no significant difference from NHA (the 95% CI of the unpaired bootstrap difference includes 0; the key gives counts per design). Error bars, 95% CI (connected-component bootstrap, 10,000 replicates). Dashed lines, prevalence.

To account for these differences, we subsampled and/or reweighted the K562 pairs to match the astrocyte screen in each property or a combination of them (Methods), and tested whether model performance on the astrocyte-matched K562 dataset became statistically indistinguishable from performance on astrocytes (Fig. 3c). We tested the matched datasets for 23 arms: 7 seq2func models with up to 3 readouts, 3 expert models, and distance as a control.

We found that enhancer–TSS distance was the main source of the difference in performance between the two cell types. Matching the distance distribution on the K562 dataset to the astrocyte dataset moved the performance of every method toward its astrocyte auPRC by a median of 55% (median over the 19 seq2func arms) and in most cases (21 out of 23; Fig. 3c) left it statistically indistinguishable from the performance on astrocytes (Supplementary Table 2). Matching target-gene expression, whose distribution differs less between the screens (Fig. 3b), had a smaller effect, with a median of 23% (14 of 23 arms no longer significantly different; Fig. 3c). However, matching distance and expression together closed a median of 75% of the gap and removed the significant difference for all 23 arms. Matching prevalence alone widened the gap for every seq2func arm by a median of 27%, because positives are more frequent in astrocytes than in K562 (5.8% against 4.5%). With prevalence matched as well, a median of 48% of the gap therefore closed (22 of 23 arms no longer significantly different; Fig. 3c).

Because the astrocyte screen contains only 133 positive pairs, these CIs are wide, and the absence of a significant difference does not imply that performance is identical. Nevertheless, benchmark composition, and enhancer–TSS distance in particular, accounts for much of the apparent drop in performance from K562 to astrocytes, suggesting that (i) seq2func models can generalize beyond K562 to a primary cell type less represented in their training data and (ii) auPRC values from different CRISPRi screens should not be compared without systematically matching their benchmark compositions.

### 4. E2GMap, a genome-wide map of enhancer–gene attribution scores across 454 biosamples, based on AlphaGenome 1

Two large-scale enhancer–gene maps are currently available: the ENCODE map based on ENCODE-rE2G’s predictions (369 cell types, tissues, and cell lines, i.e. biosamples) [9] and the recently released AlphaGenome 2 map (640 conditions covering 252 primary cell types) [25]. The ENCODE map includes biosamples for which ENCODE chromatin state data (either DNase-seq or ATAC-seq) are available, while the AlphaGenome 2 map is focused on primary cell types with scRNA-seq data available. Notably the AlphaGenome 2 map uses a model whose weights are not released, so it cannot be regenerated or extended to other elements, genes, or readouts.

We built a genome-wide enhancer–gene map for 454 biosamples, complementary to the ENCODE and AlphaGenome 2 maps. The E2GMap resource utilizes AlphaGenome 1, an open-weight model with publicly available source code, which performed as well as or better than the expert models on both screens and was not significantly outperformed by AlphaGenome 2 (Fig. 2; Supplementary Fig. 13d,e; Supplementary Table 2). AlphaGenome 1 also had higher precision than rE2G-DNase, the model behind the ENCODE map: at 70% recall (RNA-seq@exons), 0.81 against 0.51 in K562 and 0.43 against 0.20 in astrocytes (Supplementary Fig. 15a; Supplementary Table 2). Input gradients also make a genome-wide map of enhancer–gene interactions tractable, because one backward pass attributes a gene’s readout to every position in its 1-Mb window.

For each of the 20,094 protein-coding genes in GENCODE v50 [45], we centered the 1-Mb input on the gene midpoint and computed input gradients of the CAGE@every TSS and RNA-seq@exons readouts under the matched protocol (Methods). In the K562/astrocyte benchmarks we averaged the tracks of a cell type; here we attributed each of 669 tracks separately (396 RNA-seq and 273 CAGE). These tracks cover 454 biosamples (cell types, tissues, and cell lines), 95 of which have both assays. E2GScore computed one attribution value per 128 bp, which gave 8,192 positions per gene and track and 110 billion values in total (Fig. 4a). Because the map is defined by position and not by a fixed set of elements, any set of candidate elements can be scored against every gene whose window contains them.

**Fig. 4.**
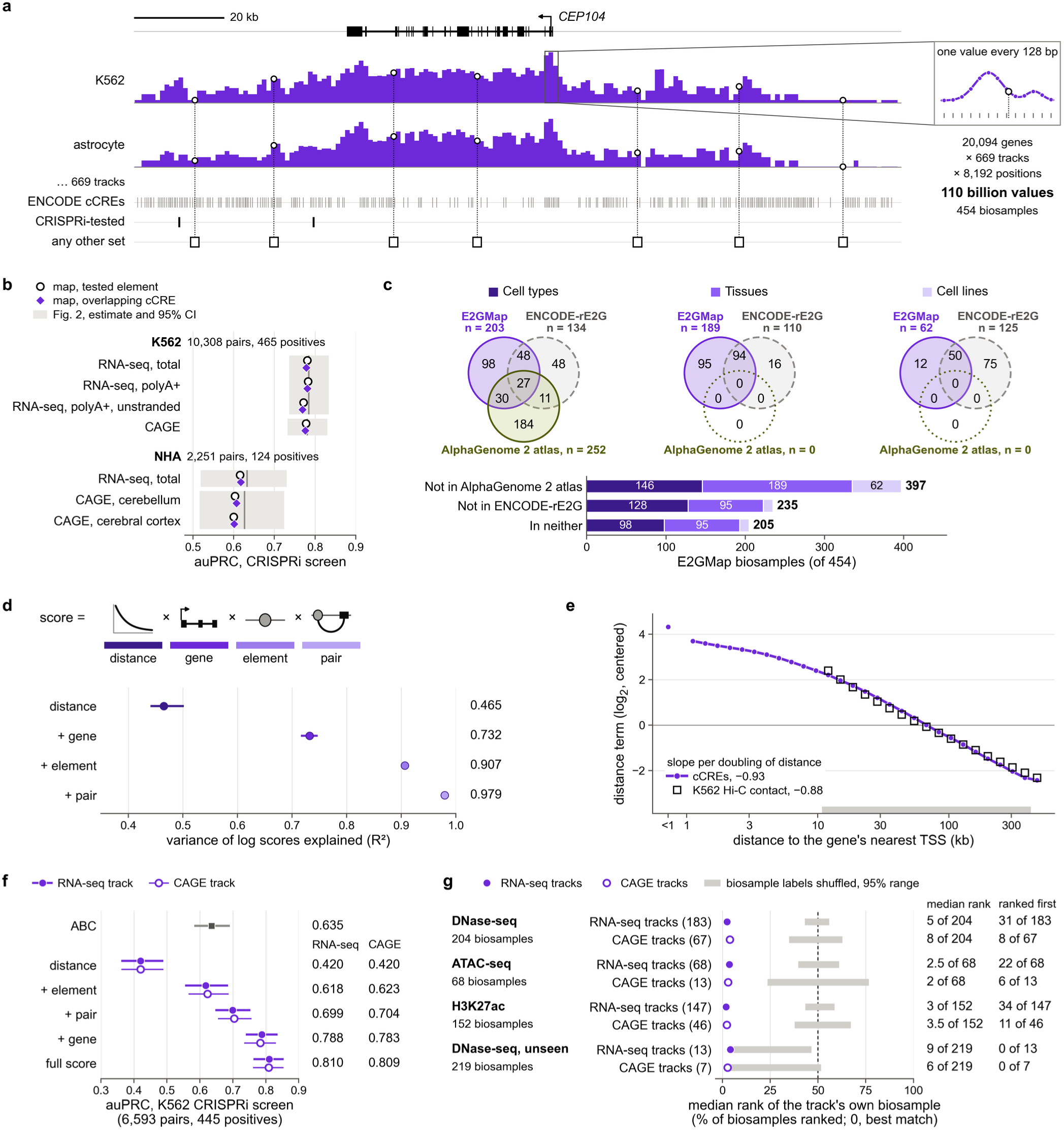
E2GMap, a genome-wide enhancer–gene attribution map, and the decomposition of its scores. **a**, For each of 20,094 proteincoding genes and 669 AlphaGenome 1 tracks, E2GMap stores input gradients over the gene’s 1-Mb window, smoothed (σ = 300 bp) and sampled every 128 bp. Profiles, 170 kb around *CEP104* in the K562 and astrocyte RNA-seq tracks (maximum per 1,024 bp, log scale). Any element set reads its scores at element midpoints (last row and inset, schematic). **b**, auPRC of single tracks of the map on the two CRISPRi screens (pairs whose gene is in the map): circles, tested element; diamonds, highest-scoring overlapping cCRE. Gray, estimate and 95% CI of the track-averaged readout in Fig. 2 (all pairs). **c**, Biosamples (ontology terms) shared by E2GMap, ENCODE-rE2G (369 terms), and the AlphaGenome 2 atlas (252 cell types across 640 conditions), by type (circles not to scale; dotted, none), and E2GMap biosamples absent from the other maps. **d**, Terms of the decomposition (Methods) and variance of log scores explained as they are added, at the 21,988,155 cCRE–gene pairs; medians and interquartile ranges over tracks. **e**, Distance term of the decomposition and of the same model fitted genome-wide to K562 Hi-C contact (16.8 million pairs), centered over the slope-fit range (gray bar, 10.8–422 kb); slopes, log₂ units per doubling of distance. **f**, auPRC on the 6,593 K562 pairs whose tested element overlaps a cCRE, as the terms are summed, and for ABC; bars, 95% CIs. **g**, Median rank of each track’s own biosample when the track-specific part of its element term is correlated across cCREs with every biosample’s chromatin peaks; gray, 95% range over 1,000 label shuffles; dashed line, chance. Unseen, DNase-seq released after AlphaGenome 1’s training data, for biosamples without AlphaGenome 1 accessibility tracks.

As a reference set, we scored the ENCODE4 Registry of candidate cis-regulatory elements (cCREs) [46]. Of its 2,348,854 cCREs, 2,198,439 (93.6%) lay in the window of at least one gene. This gave 21,988,155 cCRE–gene pairs, an average of 10.0 genes per cCRE and 1,094 cCREs per gene, and 14.7 billion scores across the 669 tracks, each available unsigned and signed.

We first checked that the map reproduces the benchmark. Scores read from single K562 and astrocyte tracks at the CRISPRi-tested elements gave auPRCs within the 95% CIs of the track-averaged AlphaGenome 1 results in Fig. 2, for both readouts and both screens (Fig. 4b). Replacing each tested element with the cCRE that overlaps it gave almost identical values (auPRC changed by −0.002 to +0.004), so the cCRE scores retain the accuracy measured in the benchmark.

To call discrete enhancer–gene links, we set one cutoff per assay: the score at which a single K562 track (total RNA-seq or CAGE) recovers 70% of K562 CRISPRi positives (Methods). On the tested pairs, the RNA-seq cutoff had a precision of 0.80 (95% CI 0.75–0.84) in K562 and, in astrocytes, which were used as a held-out evaluation set, 0.69 (0.57–0.78) at a recall of 0.53 (0.41–0.64); precision and recall on astrocytes did not differ significantly from AlphaGenome 2’s, for either its RNA-seq or single-cell expression readout (Supplementary Fig. 15b). At the *NET1* and *DNAJB6* genes, cCREs at validated enhancers passed the cutoff only in the validating screen’s cell type (Supplementary Fig. 16).

To compare the cell-type coverage of the three maps, we collapsed their biosamples to ontology terms (Methods). E2GMap’s 454 biosamples comprise 203 cell types, 189 tissues, and 62 cell lines, whereas the AlphaGenome 2 atlas covers 252 cell types, but no tissue-level conditions or cell lines; the ENCODE map covers 134 cell types, 110 tissues, and 125 cell lines (Fig. 4c). The AlphaGenome 2 atlas resolves primary cell types within tissues, including glial, retinal, and immune subtypes absent from E2GMap, while E2GMap extends coverage to bulk tissues, organs, and cell lines that the AlphaGenome 2 atlas lacks, such as endocrine and reproductive organs, the gastrointestinal tract, vasculature and peripheral nerve, and bone and skin cell types. 205 (45%) of E2GMap’s 454 biosamples are represented in neither of the other two maps: individually, 397 (87%) are not in the AlphaGenome 2 map and 235 (52%) not in the ENCODE map. Because it is built from public model weights, E2GMap can also be regenerated and extended to further biosamples, elements, and readouts.

We next asked what the E2GMap attribution scores encode. We decomposed the logarithm of the unsigned scores (14.7 billion element–gene–track values) as the sum of four categorical terms: (i) the element–nearest TSS distance, in 30 bins; (ii) an element term that varies by track (element–track; 1.5 billion levels); (iii) a gene term that varies by track (gene–track; 13.4 million levels); and (iv) an element–gene pair term (22.0 million levels). The distance and pair terms are shared by all tracks (Fig. 4d; Methods). Fitted by least squares, this factor decomposition explained about 98% of the variance of the log scores. Distance accounted for about 47%, the gene term for 27%, the element term for 17%, and the pair term for 7% (Fig. 4d). The distance term decayed similarly to Hi-C chromatin contact (Fig. 4e), suggesting that it captures similar effects on enhancer– gene regulation.

We also assessed the contribution of each term to the predictive accuracy. On the set of 6,593 K562 pairs (445 positives) whose tested element lies inside the gene’s window and overlaps a cCRE paired with that gene, the distance kernel alone gave an auPRC of 0.42. Adding the element term raised it to 0.62, adding the pair term gave 0.70, and adding the gene term 0.79, close to the 0.81 of the full score; RNA-seq and CAGE tracks behaved similarly (Fig. 4f).

The element term isolates the part of each score that depends on the cCRE and the track but not on the gene. Because AlphaGenome learns chromatin and expression from the same sequence features, we expected this term to capture the chromatin activity of cCREs in the track’s biosample, even though E2GMap was built from expression readouts alone. To test this, we correlated each track’s element coefficients across cCREs with the DNase-seq, ATAC-seq, and H3K27ac ChIP-seq signal of every biosample that was available (Methods). The correlation with the track’s own biosample (median r = 0.13 for DNase-seq, 0.18 for ATAC-seq, and 0.15 for H3K27ac in RNA-seq tracks) was higher than with nearly all other biosamples: in RNA-seq its median rank was 5th of 204 for DNase-seq, 2.5 of 68 for ATAC-seq, and 3rd of 152 for H3K27ac; CAGE had similar ranks (Fig. 4g). For the 13 RNA-seq tracks whose biosamples had DNase-seq data that were not used to train AlphaGenome, the result was similar (median rank 9th of 219). The agreement therefore does not depend on the model having been trained on that biosample’s chromatin data.

E2GMap, comprising the cCRE–gene scores and the fitted terms, is available at e2gscore.com.

## Discussion

Seq2func models are increasingly used to infer enhancer–gene regulation, although they are trained to predict molecular readouts rather than functional regulatory interactions. As successive models have introduced longer sequence contexts and more sophisticated architectures, their enhancer–gene predictions have also been evaluated using different attribution procedures [22–24]. Here, we introduced E2GScore, a unified framework for benchmarking enhancer–gene predictions from seq2func models, and showed that recent models accurately identify regulatory interactions across cell types, while apparent differences in performance are strongly influenced by benchmark composition. We further introduced E2GMap, a genome-wide map spanning 454 biosamples, and decomposed its attribution scores to investigate the contributions of enhancer–gene distance, regulatory element activity, gene-specific properties, and enhancer–gene specificity.

Our findings highlight the importance of distinguishing improvements in the underlying models from those introduced by the scoring procedure. E2GScore provides a reproducible framework for making this distinction, while establishing practical choices for extracting enhancer–gene predictions that are both accurate and computationally efficient.

Our results also demonstrated substantial progress in the ability of seq2func models to identify distal regulatory interactions. Earlier studies found that Enformer captured promoter-proximal regulatory effects but largely underestimated the contribution of distal enhancers to gene expression [28]. More recent models show markedly improved discrimination of functional distal enhancer–gene pairs, with AlphaGenome performing as well as or better than supervised predictors that rely on experimentally measured chromatin features and CRISPRi training data [5,9,10]. This is particularly encouraging for cell types in which the experimental data required by supervised models are unavailable. However, the ability to discriminate functional from non-functional enhancer–gene pairs does not necessarily imply accurate prediction of the magnitude or direction of regulatory effects [31], an important limitation for interpreting non-coding variants.

Previous studies have highlighted how benchmark construction can inflate enhancer–promoter prediction accuracy [36,37]. We extend this concern to comparisons across CRISPRi screens, showing that differences in enhancer–TSS distance and target-gene expression can account for much of the apparent loss of performance in astrocytes relative to K562 cells. These findings caution against interpreting absolute auPRC differences between screens as evidence of poorer generalization ability and argue for evaluating models on benchmarks matched for relevant properties. Although our results support substantial transfer of predictive performance to primary cells, the limited number of positive astrocyte pairs leaves uncertainty about the magnitude of any remaining cell-type-specific differences.

The development of AlphaGenome 2 introduces an important new capability: conditioning sequence-based predictions on primary cell identities and states represented by single-cell transcriptomic data [25]. In our evaluation, where we estimated uncertainty with CIs, we did not observe a statistically significant improvement in enhancer–gene linking over AlphaGenome 1 for either K562 or astrocytes. These results may be due to insufficient power to detect moderate differences with the current size of CRISPRi screens, but suggest that the benefits of cell-state conditioning for predicting functional enhancer–gene interactions remain to be established. Larger perturbation datasets spanning diverse primary cell types and cellular states will be needed to determine whether this additional modeling complexity translates into improved enhancer–gene prediction.

Finally, E2GMap extends sequence-derived enhancer–gene predictions to cellular contexts that complement existing atlases [9,25,47]. Its construction from publicly available model weights and position-resolved attribution scores also allows regulatory links to be re-evaluated using alternative candidate elements and annotations. More fundamentally, the decomposition of attribution scores offers insight into the regulatory information learned by seq2func models.

Together, E2GScore and E2GMap provide a framework for evaluating, mapping, and interpreting enhancer– gene regulation from DNA sequence. Our findings demonstrate the importance of standardized scoring procedures and carefully matched benchmarking datasets for assessing advances in seq2func models. The size of current CRISPRi screens limits how finely models can be separated, underscoring the importance of larger perturbation datasets across diverse cell types. Nonetheless, our results highlight the growing ability of seq2func models to predict functional regulatory interactions across cell types, while E2GMap provides an extensible resource that complements existing genome-wide enhancer–gene maps.

## Methods

### E2GScore framework

E2GScore separates the scoring algorithm (42 parameters) from the model. A single experiment, which scores one model on one CRISPRi screen under one protocol, requires: (1) the scoring algorithm, (2) a backend module exposing the model’s forward pass and input gradient, (3) the model’s published weights, (4) a YAML configuration file setting the protocol’s parameters, (5) the screen’s table of candidate enhancer–gene pairs, (6) GENCODE exon and/or TSS annotations (v50 in our protocol), and (7) the hg38 reference genome. Adding a new model requires a backend module and the model’s geometry and track metadata, and does not change the algorithm. The algorithm exchanges only NumPy arrays with a backend, so a backend can in principle use any framework, provided it can differentiate the gene readout. Each run writes a table of per-pair scores.

We also optimized E2GScore for speed. For input gradients, one forward and backward pass per gene, replicate, and augmentation scores all candidate enhancers in the context window. For AlphaGenome, the decoder and output head run only over the span of the readout positions plus a 2-kb halo. We also provide a cached ISM path, similar to fastISM [48], which, on samples of the astrocyte CAGE@TSS runs, changed no pair’s rank (largest score difference 3.2 × 10⁻⁵ of the 99th-percentile score) and ran 2.1–5.2 times faster (Enformer, 3.2; Borzoi, 2.1; Flashzoi, 2.1; AlphaGenome 1, 5.2). The reference sequence is encoded once per gene, replicate, and augmentation; for each of the N shuffles, only the shuffled window ±4 kb is re-encoded and spliced in, and the transformer and decoder are re-run. This is possible because convolutions and pooling give the encoder a finite receptive field, and its normalizations use fixed statistics.

### Reproducing published protocols

We reproduced Fig. 2b and Extended Data (ED) Fig. 7c of Avsec et al. [22]; Fig. 4c,d, the flat-window comparison, and Supplementary Fig. 7b of Linder et al. [23]; Fig. 4j and ED Fig. 7a of Avsec et al. [24]; and Fig. 4b,c and Supplementary Fig. 11a–c,f of Gayoso et al. [25], each for the paper’s own model. The E2GScore framework scored every arm of the first three papers; only its parameters changed: the protocol, the attribution method (input gradients, ISM, attention), the model (Enformer, Borzoi, and AlphaGenome, as the PyTorch ports lucidrains/enformer-pytorch, johahi/borzoi-pytorch, and genomicsxai/alphagenome-pytorch [49]), and the screen [5,6,8,9]. AlphaGenome 2, whose weights are not released, was scored through the AlphaGenome API with DeepMind’s own scoring function, which on identical attributions agrees with E2GScore’s to within 4 × 10^−16^ except at N bases. auPRC was computed as step-wise average precision, the estimator of Linder et al. [23], Avsec et al. [24], and Gayoso et al. [25]; Avsec et al. [22] do not state theirs. A published auPRC was counted as reproduced when it lay inside our 95% percentile interval over 10,000 bootstrap resamples of the pairs of its distance bin.

Because Avsec et al. [22] did not release their enhancer–gene pairs, we rebuilt the Fulco (Fulco et al. [5], their Supplementary Table 3a) and Gasperini (GEO GSE120861) cohorts in hg38; their per-distance-bin pair counts matched the published counts to 98.2% and 98.3%. We digitized the published auPRC values, which are given only as plots. The 196,608-bp windows were centered on the major TSS, the GENCODE v32 TSS with the highest predicted K562 CAGE. The readout was CAGE summed over the three 128-bp bins at that TSS (the two K562 tracks; all 638 CAGE tracks for ED Fig. 7c), with no gradient correction or normalization. Input gradients were summed over the 2 kb centered on the enhancer, with scores cropped to ±57,344 bp of the TSS as in DeepMind’s Enformer notebook. ISM replaced the 2 kb with one random sequence (a segment of the same input window for K562, random bases for ED Fig. 7c) and scored |f(modified) − f(reference)|. Attention from the three TSS tokens, averaged over heads and layers, was summed over the 16 tokens (2,048 bp) centered on the enhancer midpoint. Pairs were binned as in their Fig. 2b (to 131 kb) and none were dropped: pairs outside the input window, or outside the crop for gradients, scored 0.

For Linder et al. [23], we scored the authors’ FlowFISH+ [8] and Gasperini pair tables and annotation (GENCODE v38 basic protein-coding; the paper states v41), obtained on request, against the values in their notebooks. The 524,288-bp windows were centered on the gene midpoint (Calico’s merged-exon centroid). The readout was the log of K562 RNA-seq summed over exon bins, on Borzoi’s uncompressed scale and averaged over the gene-strand half of the 20 stranded tracks. We applied gradient correction over the four bases, averaged as signed maps over both orientations and the four replicates, and divided by the mean absolute input gradient. Each pair was then scored with a Gaussian (σ = 300 bp, ±1,200 bp) or a flat 2-kb mean. As in the released code, ISM permuted the 1,024 non-overlapping dinucleotides of the 2,048 bp at the enhancer (a 2-mer block shuffle) 16 times per orientation and replicate, and scored (1/128) Σ [log f(reference) − log f(modified)] over the 128 passes, keeping its sign. Pairs outside the window were dropped.

For Avsec et al. [24], we scored the released ENCODE-rE2G K562 benchmark [9], restricted to genes in GENCODE v46 (10,353 pairs, 471 positives). The 1,048,576-bp windows were centered on the gene midpoint; the readout and gradients were the same as for Linder et al. [23], with one model and no reverse-complement augmentation. For the RNA-seq variant-effect (ISM) bar, the 2 kb at the enhancer was permuted 10 times via a mononucleotide shuffle and scored |(1/10) Σ [log f(modified) − log f(reference)]|. Enhancers outside the window scored 0.

For Gayoso et al. [25], whose AlphaGenome 2 weights are not released, we scored three benchmarks through the AlphaGenome API (client 2.0.0) on the paper’s own pairs. These were the ENCODE-rE2G K562 benchmark restricted to genes in GENCODE v46 [9] (10,353 pairs, 471 positives), the evaluation subset of the astrocyte screen [10] (1,902 pairs, 122 positives), and the ENCODE-rE2G held-out benchmark [9] (screens in K562, WTC11, HCT116, GM12878, and Jurkat cells; 4,378 pairs, 190 positives). The API fixes the window: 1,048,576 bp centered on the median TSS of the gene’s GENCODE v46 transcripts. Each gene took one input-gradient call, conditioned on K562 (EFO:0002067), brain astrocyte (UBERON:0000955_CL:0000127), or the pair’s own cell line, with observed tracks only. Its objective was the log of the gene’s predicted expression averaged over those tracks: bulk RNA-seq over exons for the cell lines and single-cell gene expression for astrocytes, as in the paper. Each element was scored with the client’s score_enhancers function. It takes the gradient corrected over the four bases, in absolute value, averaged with a Gaussian (σ = 300 bp) over the 2,400 bp around the element midpoint. For the window-normalized variant, the score was divided by the mean absolute input gradient over the window. Elements outside the window scored 0. Following the authors, held-out pairs were weighted by the benchmark’s probability of a direct effect. Distances were those that reproduce the paper’s 1/distance auPRC (within 0.0003 in every bin): each benchmark’s own distance column for the astrocyte and held-out benchmarks, and for K562 the TSS distance in DeepMind’s released K562 prediction file. Published values were read from the paper’s bars, since it prints few of them.

### Preprocessing CRISPRi screens

The K562 dataset [9] combines three CRISPR screens (Nasser et al. [8], mostly CRISPRi-FlowFISH; Gasperini et al. [6], Perturb-seq; Schraivogel et al. [7], TAP-seq), harmonized by the authors, and excludes elements that overlap an annotated promoter or the target gene’s body. Of its 10,356 pairs, 471 are positives (a significant decrease in expression, adjusted P < 0.05) and 9,885 are negatives, which showed no significant decrease despite at least 80% power to detect a 15% decrease (25% for the CRISPRi-FlowFISH data).

The NHA screen [10] tested 7,759 pairs and called 158 hits (Benjamini–Hochberg false discovery rate (FDR) < 0.1 on empirical P values from negative-control guides), 133 with decreased and 25 with increased expression. We used the modeling table of Green et al. [10], which keeps a pair if it has at least 80% power to detect a 15% decrease in expression (i.e. well-powered pairs) or if it is a hit with decreased expression. This gives 2,307 pairs comprising 133 positives (the downregulated hits) and 2,174 negatives (2,168 well-powered non-hits and the 6 well-powered upregulated hits).

### Varying key protocol parameters

For input gradients, we varied cross-gene normalization via two parameters: log_scalar (default “true”) and normalise (“none” in our matched protocols; the code default is “global”). Gene-readout normalization (log_scalar: true) applies a logarithmic transform to the predicted gene readout *u* (here *u* is the summed readout *S* of equation (1)) before backpropagation, yielding the gradient map d log(*u*)/d*x* = (1/*u*) d*u*/d*x*, i.e. the gradient map d*u*/d*x* divided by the positive per-gene expression constant *u*. Gradient-map normalization (normalise: global) (what Gayoso et al. [25] call “window-normalization”) divides the enhancer–gene score by the mean absolute input gradient mean(*|*d*u*/d*x · x|*). Combining both (as Linder et al. [23] and Avsec et al. [24] do) is equivalent to gradient-map normalization alone, making the logarithmic transform redundant. This is because gene-readout normalization divides the map *M* by the gene read-out *u*, and gradient-map normalization then divides by the mean of that same quantity, so *u* cancels: *s*(*e, g*) = *|M* /*u|*/ mean(*|M* /*u|*) = *|M |*/ mean*|M |*.

For ISM, we tested cross-gene normalization by varying the ism_statistic (code default “abs_mean_diff”) parameter (the benchmarks report “log_abs_diff”, equation (4), computed from the stored shuffles). For no normalization, we used the absolute difference between the perturbed and wild-type gene readouts averaged over shuffles (ism_statistic: abs_diff). We took a log fold-change (ism_statistic: log_abs_diff) of the gene readout as well as a relative change (ism_statistic: rel_abs_diff).

To test the importance of cell-type specificity we varied the cell_type parameter. We selected the K562 or astrocyte CAGE tracks (cell_type: k562, astrocytes) predicted by each model for the readout u, as well as all CAGE tracks (cell_type: all) to represent the cell-type-agnostic readout. We randomly sampled a matched number of tracks from each model’s set of non-K562 CAGE tracks 10 times without replacement (cell_type: rand2-s0–s9 for Enformer and Borzoi, rand1-s0–s9 for AlphaGenome) to control for the differing number of tracks between these two groups.

We varied the TSS used by the CAGE protocol (anchor: single-tss; assay: cage); the chosen TSS sets both the center of the input window and the bins from which the CAGE signal is read. We tested each screen’s provided TSS: for K562 (Gschwind et al. [9], the distanceToTSS column of the released benchmark table) this was the RefSeq TSS with the most associated coding transcripts, used by ABC and ENCODE-rE2G, while for NHA (Green et al. [10], the modeling table of the authors’ repository) it was the 5′-most TSS (GENCODE v32). We also tested the TSS for the MANE Select transcript. Lastly we tested the TSS with the highest predicted CAGE signal in the cell type of interest (“major predicted TSS”), and the TSS with the highest experimentally measured CAGE signal in the cell type of interest (“major measured TSS”).

We varied the number of gene readout bins for the CAGE protocol via the tss_width parameter: 128, 384, and 640 bp. To test whether the gene readout was shift-invariant to the choice of input window anchor or whether certain anchors were favorable to CAGE/RNA-seq protocols, we varied the anchor parameter: single-tss [22], using the MANE Select transcript’s TSS, or gene-midpoint, the mean position of the gene’s exonic bases [23]. We combined this choice of anchor with the CAGE (region: tss; assay: cage) or RNA-seq (region: exons; assay: rnaseq) protocol. Lastly, we tested with and without the gradient correction step from Majdandzic et al. [50], varying the mean_subtract parameter (true/false); varied the shape of the Gaussian kernel to aggregate scores at the enhancer loci: 12 kernels (gaussian_sigma 147–1,605 bp, gaussian_truncate 1–3 standard deviations; matched in second moment to flat windows of 500, 1,000, 2,000, and 3,000 bp), each compared with the protocol kernel (300 bp, truncated at 4 standard deviations); and compared average pooling (enh_agg: mean) to Gaussian pooling (enh_agg: gaussian) of the same second moment at fixed widths (enh_span): 500, 1,000, 2,000, and 3,000 bp.

### Matched protocols across sequence-to-function models

Besides Enformer, Borzoi, and AlphaGenome, we scored three further models with the same algorithm: Flashzoi, Cerberus, and NTv3. We used E2GScore to score an enhancer–gene pair by attributing the model’s prediction for the gene’s expression to the candidate enhancer, using input gradients or ISM. The input *x* is the one-hot hg38 sequence (*L ×* 4, where *L* is the model’s input length), centered on an anchor. The gene readout is

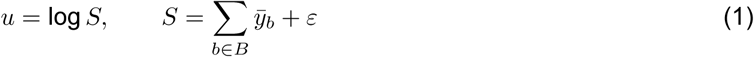

where *B* is a set of output bins, *y̅_b_* is the prediction in bin *b* averaged over the selected tracks, and *ε* = 10^−6^. The logarithm is natural for Enformer, Borzoi, Flashzoi, and Cerberus, and base 2 for NTv3 and AlphaGenome.

#### Readouts and anchors

CAGE@TSS reads CAGE over the 384 bp centered on the gene’s TSS (3 bins for Enformer, 12 for Borzoi, Flashzoi, and Cerberus, and 384 for NTv3 and AlphaGenome). The input is centered on that TSS, as in Avsec et al. [22]. We used the MANE Select TSS (for genes without one, the Ensembl canonical TSS, or else the screen’s TSS). CAGE@every TSS reads CAGE over the union of promoters, i.e. 384-bp windows centered on the 5′ ends of all the gene’s GENCODE v50 transcripts, merging any overlapping windows into a single promoter. RNA-seq@exons (all models except Enformer) reads RNA-seq over every bin that has any overlap with an exon of these transcripts, merging any overlapping exon bins. For both, the input is centered on the gene midpoint. RNA-seq@exons with this anchor follows Linder et al. [23] and Avsec et al. [24]. Bins outside the region the model predicts were excluded. Enformer predicts only the central 114,688 bp of its input, and NTv3 only the central 393,216 bp; gradients still cover the whole input, so enhancers anywhere in the window can be scored. A gene with no TSS in the predicted region cannot be read, so its pairs score 0, as do pairs whose enhancer lies outside the window.

#### Tracks

For K562, we read K562 CAGE and RNA-seq tracks. For NHA, we read CAGE from primary astrocytes of the cerebral cortex and cerebellum, and astrocyte RNA-seq (Supplementary Table 3). Stranded tracks were read on the gene’s sense strand. For Borzoi and Flashzoi we inverted the target compression [23], and for Cerberus and NTv3 each model’s own output transform, before summing bins, and computed u for each replicate model (four for Borzoi and Flashzoi, eight for Cerberus, and one for NTv3).

#### Input gradients

One backward pass per gene (per replicate model for Borzoi, Flashzoi, and Cerberus) gives *∂u*/*∂x* and scores every enhancer in the window at once. We corrected the gradient and read it at the reference base, which equals the corrected gradient times the input:

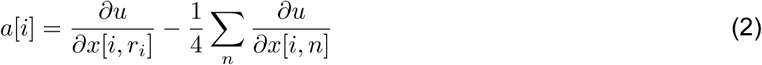

where *r_i_* is the reference base at position *i* and *n* ∈ {A, C, G, T}. For Borzoi, Flashzoi, and Cerberus, the replicates’ signed maps were averaged before taking absolute values [23]. The score is the Gaussian-weighted mean of *|a|* around the enhancer midpoint *c_e_*, using the kernel of Linder et al. [23] and Avsec et al. [24]:

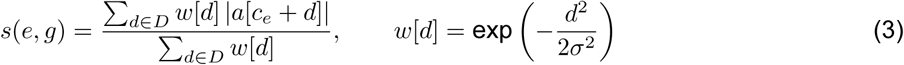

where *σ* = 300 bp and *D* holds the offsets *|d| ≤* 1,200 bp (4*σ*) for which *c_e_* + *d* lies inside the window. Pairs whose enhancer midpoint lay outside the window scored 0.

#### Normalization

We used only gene-readout normalization (equation (1)). Because *u* is a logarithm, *∂u*/*∂x* ∝ (*∂S*/*∂x*)/*S*, so each gene’s map is divided by its own predicted signal distributed among promoter or exon bins. For CAGE@every TSS and RNA-seq@exons, each promoter or exon is weighted in proportion to its share of the total predicted CAGE or RNA-seq coverage over bins so as to account for silent and dominant promoters. Setting *T* = ∑*_b_T_b_*:

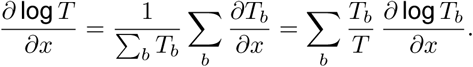

In the same way, each averaged track is weighted by its predicted signal. A silent promoter or exon therefore contributes little to the gene’s map, and a dominant one most of it.

#### In silico mutagenesis

For each enhancer, we replaced the 2,000 bp centered on its midpoint (clipped at the window edge) with a dinucleotide-preserving shuffle [51]. We drew *N* = 10 shuffles *x̃*_1_*, …, x̃_N_* and scored

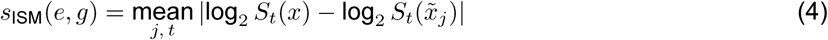

where *S_t_*=∑*_b∈B_ y_b,t_* is the readout of track *t* alone. The mean runs over shuffles, tracks, and, for Borzoi, Flashzoi, and Cerberus, replicates. ISM thus costs *N* forward passes per enhancer and replicate model, plus one reference pass per gene.

AlphaGenome 2 [25], whose weights are not released, was scored through the AlphaGenome API as for its reproduction: one input gradient call per gene with the K562 or brain astrocyte condition, for the bulk RNA-seq readout over exons and for the single-cell gene-expression readout, in a 1-Mb window that the API centers on the median TSS of the gene’s transcripts; elements were scored with the client’s score_enhancers function, without window normalization.

### Expert models

We ran the ENCODE-rE2G logistic regression models (DNase and extended versions) using the model training code and the input features provided by Gschwind et al. [9], their Supplementary Tables 3 and 8. Similarly we ran the EGrf random forest models using our Python port of the model training code (in R) and the input features provided by Green et al. [10], their Supplementary Table 9.

The ABC score was defined according to Fulco et al. [5] as (*A_e_* x *C_e,p_*)/(∑*_e′_A_e′_* x *C_e′,p_*) where *A_e_* = 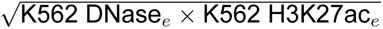 and *C_e,p_* = K562 Hi-C*_e,p_*. Here *e* is the candidate enhancer, *p* is the promoter, and *e^′^* is any candidate enhancer within 5 Mb of the gene’s chosen TSS associated with *p*. ABC scores were taken as released, not recomputed: for K562 from the feature table released by Gschwind et al. [9], and for NHA from the ENCODE astrocyte predictions (ENCFF440FMQ), a DNase-only ABC score with cell-type-averaged Hi-C, to which the K562 inputs above do not apply. Enhancer–TSS distance was defined as the absolute distance between the MANE Select transcript’s TSS and the enhancer midpoint.

For ENCODE-rE2G and EGrf models, we obtained predictions via hold-one-chromosome-out cross-validation: a model was trained on 22 chromosomes and performed inference on the held-out chromosome; we cycled through all 23 chromosomes as the held-out chromosome and concatenated all held-out predictions, following the procedure of Gschwind et al. [9]. EGrf, which Green et al. [10] trained with random folds, was refitted under the same scheme, so that every supervised model is evaluated out of fold in the same way. The predictions published by Green et al. [10] were also scored on the same pairs, as a check (Supplementary Table 2). rE2G-extended and EGrf-extended were evaluated in K562 only, because their additional features come from K562 assays not available for astrocytes (such as EP300 ChIP-seq and ChIA-PET).

### Benchmarking models

We report each method’s auPRC with a 95% CI. When comparing two methods, we report the paired difference in auPRC (ΔauPRC), its 95% CI, and a two-sided bootstrap P value for H_0_: ΔauPRC = 0. Pairs that share an enhancer or a gene are not independent, so we resampled the connected components of the bipartite enhancer–gene graph formed by each screen’s tested pairs (663 in K562, 309 in NHA). Each of 10,000 replicates draws as many components as the screen has, with replacement, and keeps every pair of each drawn component; all methods are evaluated on the same replicates. CIs are the 2.5th and 97.5th percentiles of the replicates and are not adjusted for multiple comparisons. The P value is twice the smaller of the fractions of replicate ΔauPRCs at or below 0 and at or above 0 (at most 1); with 10,000 replicates its resolution is 2×10^−4^, and comparisons with no replicate on the other side of 0 are reported as *P <* 2 × 10^−4^. Differences between the two screens (K562 − NHA) for one method are unpaired: their two-sided P value was computed over all pairs of the two screens’ bootstrap replicates.

We controlled the false discovery rate with the Benjamini–Hochberg procedure, using one family per figure and screen: all comparisons drawn in a figure, together with that figure’s rows in Supplementary Table 2, were adjusted together, separately for K562 and NHA, and a figure’s between-screen differences (K562 minus NHA) formed a family of their own. A comparison over all pairs and the same comparison in each distance bin are separate tests within one family. We call a comparison significant at q < 0.05. Supplementary Table 2 lists each comparison with both auPRCs, ΔauPRC, its 95% CI, the P value, the q value, and its family, and its Read me sheet gives the size of every family.

auPRC is computed as tie-aware step-wise average precision (AP). A method scores 0 on pairs it cannot score, so its all-pairs AP mixes how it ranks pairs with how many pairs it can score. A seq2func model can score a pair when the element midpoint lies inside its input window centered on the anchor. ABC has no score (0) for 2,732 K562 pairs, all without Hi-C contact in the release, and for the 405 NHA pairs without an ENCODE astrocyte prediction; rE2G-DNase in NHA, whose features come from the same predictions, is 0 on the same 405 pairs. rE2G in K562 and EGrf score every pair (EGrf’s zeros are predictions).

#### Reach and ranking

A seq2func model can score a pair if the enhancer lies inside its input window and, for CAGE@every TSS, at least one annotated TSS of the gene lies inside the region the model predicts; ABC scores pairs with a score above 0, rE2G-DNase scores pairs with a non-zero score (every pair in K562), and rE2G-extended, EGrf, EGrf-extended, and 1/distance every pair, as in Fig. 2. Pairs a method cannot score take its lowest score and tie at the bottom. AP (auPRC) is tie-aware, as in Fig. 2 (tied pairs enter the precision–recall curve together); the identities below are exact under this convention.

Let *N* and *P* be the numbers of pairs and positives. For a set of pairs *C* with *P_C_* positives, let AP*_C_*(*X*) be the AP of method *X* on *C* alone and *X|C* its scores with every pair outside *C* set to −1 (all scores are ≥ 0). Then

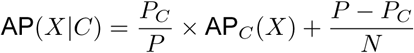

For one method, take *C* as the pairs *X* can score (so *X|C* ranks as *X* does). Its reach is *w* = *P_C_*/*P*, AP(*X*) = *w ×* AP*_C_*(*X*) + (1 *− w*) *× P* /*N*, and its reach penalty, the auPRC lost to the tie at the bottom, is

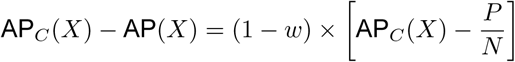

Methods that score every pair (*w* = 1) have no penalty and are omitted from Supplementary Fig. 11. For two methods *A* and *B*, take *C* as the pairs both can score; the auPRC gap is then

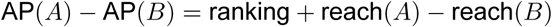

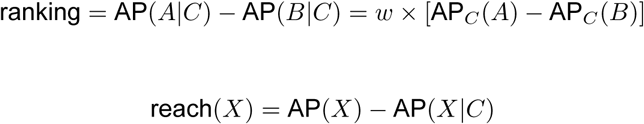

Ranking is the difference on the shared pairs, weighted by their share of positives; reach(*X*) is the signed contribution of the pairs only *X* can score (0 if there are none). We compared AlphaGenome 1, which unlike AlphaGenome 2 has all three readouts, with each Enformer, Borzoi, Flashzoi, NTv3, and Cerberus arm under the same readout (14 comparisons per screen); AlphaGenome 2 scores almost exactly the same pairs (one positive fewer in K562), so its reach terms would barely differ. reach(*B*) was 0 in every comparison, so Supplementary Fig. 12 shows the reach of AlphaGenome 1 and its share of the gap, reach(*A*)/[AP(*A*) *−* AP(*B*)]. All terms were computed on Fig. 2’s 10,000 bootstrap replicates (connected components of the enhancer–gene graph resampled with replacement), with 95% CIs; Supplementary Table 2 gives two-sided bootstrap P values, as for Fig. 2, and Benjamini–Hochberg q values within each screen with the ranking and the reach terms of the 14 comparisons in one family (28 tests).

### Other comparisons

Within-gene and within-enhancer areas under the receiver operating characteristic curve (auROC) are the auROC of each gene’s, or each enhancer’s, pairs, averaged over the genes or enhancers with at least one positive and one negative pair (Supplementary Fig. 13b,c), with 95% CIs from 10,000 bootstrap resamples of connected components. Precision at 70% recall is the precision at the first score threshold, in descending order, at which a method recovers at least 70% of a screen’s positives; tied pairs enter together, and pairs a method cannot score take a score of 0 (Supplementary Fig. 15a). AlphaGenome 1 (RNA-seq@exons and CAGE@every TSS) was compared with the supervised models and ABC on the bootstrap resamples used for auPRC, with Benjamini–Hochberg correction within each screen (10 tests in K562, 6 in NHA; Supplementary Table 2).

### Gene expression prediction

We predict the expression of 2,109 K562 and 1,052 NHA genes. The RNA-seq results use 2,094 and 1,038 genes because RNA-seq was read over the MANE Select coding sequence, which the 15 and 14 non-coding genes lack; Enformer’s CAGE results use 2,086 and 1,040 genes, because with the gene-midpoint anchor the others have no annotated TSS inside the 114,688 bp that Enformer predicts. CAGE was read as CAGE@every TSS, and RNA-seq as the mean, not the sum, of predicted coverage over the coding-sequence bins. Measured CAGE for K562 was taken from the models’ own training libraries, so the CAGE comparison is in-sample: K562 combines CNhs11250 (weight 1/2) with CNhs12334–6 (1/6 each), while astrocyte is the mean of the cortex and cerebellum library groups. Measured RNA-seq transcripts per million (TPM) for K562 was from ENCSR000AEM (mean of two replicates), while for NHA it was from bulk RNA-seq (mean of two dCas9-KRAB background lines) [10]. We compute Pearson correlations between log₂(1+T) values over all genes, with 95% CIs from 2,000 gene-resampling bootstrap draws.

### Matching benchmark composition

To compare enhancer–TSS distance and target-gene expression distributions between cell types we used the two-sample Kolmogorov–Smirnov (KS) test. We reported Benjamini–Hochberg-adjusted P values, obtained by permuting connected components over 10,000 shuffles, with the four per-class tests of Fig. 3a,b and the two all-pairs tests in one family (Supplementary Table 4).

We matched K562 to NHA for all models except rE2G-extended and EGrf-extended (which are only evaluated for K562) using five designs and recomputed each method’s auPRC (weighted average precision) from its existing enhancer–gene scores, without re-running any model. The designs matched: (1) total and positive counts only (N), (2) enhancer–TSS distance only, (3) target-gene expression only, (4) distance + expression, and (5) N + distance + expression. In every design, positives were matched only to positives and negatives only to negatives. Designs that matched distance or expression compared distributions in bins at NHA’s quantiles (10 for positives, 20 for negatives; for distance, quantiles of log_10_ distance). The outer edges were NHA’s minimum and maximum: 17 bp–497 kb (positives) and 2.5–588 kb (negatives) for distance to the MANE Select TSS (to the screen’s own GENCODE v32 TSS, every pair lies 2.4–500 kb away) and the 30th–99.97th percentile for expression. A K562 pair outside a matched variable’s range, or without an expression value when expression was matched, received weight 0. Weights were then rescaled so that each class kept its size. To match expression and distance variables at once, we raked the weights (iterative proportional fitting) [52]. Starting from weight 1, the weights were rescaled so that each expression bin summed to its target (the class size times NHA’s share of that class in the bin), and then the same was done for the distance bins. This cycle was repeated until no bin differed from its target by more than 10^−6^ of the class size, for at most 200 cycles. Expression was each target gene’s percentile of TPM within its own cell type, among the 19,844 protein-coding genes quantified in both cell types (TPM sources as in ‘Gene expression prediction’).

To match N, we drew NHA’s numbers of positives and negatives (133 and 2,174) from K562 uniformly at random without replacement, with 10 random seeds; distance and expression kept K562’s distributions. To match distance, we weighted each K562 pair by NHA’s share of its class in the pair’s distance bin, divided by K562’s share. To match expression, we raked all K562 pairs to NHA’s expression bins and to K562’s own distance bins (at K562’s quantiles, with K562’s shares as targets). Expression thus took NHA’s distribution while distance kept K562’s. To match distance + expression, we raked all K562 pairs to NHA’s expression and distance bins. To match N + distance + expression, we visited NHA’s pairs of each class in random order. Each NHA pair took an unused K562 pair of the same class, chosen uniformly at random from those whose log_10_ distance differed from its own by at most 0.02 (a factor of 1.047), or else the nearest unused pair. This subsample was then raked to NHA’s expression and distance bins, with 10 seeds. We verified with weighted KS distances that the matched distributions agreed well (Supplementary Table 4) and determined whether the matched auPRC was significantly different from the NHA auPRC. Each matched auPRC was compared with the NHA auPRC by an unpaired bootstrap: randomly permuted NHA replicates were subtracted from the matched K562 replicates (10,000 connected-component replicates per screen, each pair’s weight multiplied into its component’s count), and the two were called different when the 95% percentile interval of the difference excluded 0, without adjustment for multiple comparisons. The share of the gap closed by a design is (K562 auPRC − matched auPRC)/(K562 auPRC − NHA auPRC), and we report its median over the 19 seq2func arms and, per arm, its 95% percentile interval over the bootstrap replicates.

### Computing E2GMap

For each protein-coding gene in GENCODE v50 (comprehensive annotation; chromosomes 1–22, X, and Y), every cCRE of the ENCODE4 Registry (all eight classes) whose midpoint lay within the 1,048,576-bp input window was paired with the gene. We used the CAGE@every TSS and RNA-seq@exons readouts with gene-readout normalization, computing u (equation (1)) separately for each track. The 669 tracks (396 RNA-seq and 273 CAGE) come from 454 biosamples: 285 with RNA-seq, 264 with CAGE, and 95 with both. We grouped them into 563 biosample units, one per biosample and assay: 299 for RNA-seq, in which 14 tissues are split into fetal and postnatal units, and 264 for CAGE. A unit holds more than one track when its biosample has several libraries of that assay (97 extra RNA-seq tracks and 9 extra CAGE tracks). Each track retains its library protocol (polyA+ or total RNA-seq; hCAGE or LQhCAGE), strandedness, source (RNA-seq: ENCODE or GTEx; CAGE: FANTOM5) [53–55], and life stage. For each gene, one forward pass was followed by one backward pass per track. Each backward pass gave the corrected gradient (equation (2)) over the whole window, from which all the gene’s cCREs were scored with equation (3) for unsigned scores and equation (3) without the absolute value for signed scores.

We used the AlphaGenome all-folds checkpoint, which was trained on all chromosomes, through the community PyTorch port [49] (v0.1.dev1+g473d3d226) with PyTorch 2.6.0 and CUDA 12.4, in fp32 without mixed precision. To fit in 141 GB VRAM, the decoder at 1-bp resolution was evaluated only over the readout region plus 2,048 bp on either side, more than the decoder’s reach of 762 bp on each side, so readouts and gradients equal those of a full evaluation to floating-point round-off. Genes were processed in parallel, each on one NVIDIA H200 GPU on the University of New South Wales’ Katana cluster, for about 4,650 GPU-hours over 26 days. Peak memory reached about 110 GiB for the typical gene; for 258 long genes whose predicted peak exceeded 135 GiB, we used checkpointing. Windows extending past a chromosome end were padded with N, encoded as all-zero rows of the one-hot input. For 154 genes the window did not contain the whole readout region (for 68 the region is longer than the window): their readouts summed only the exons and TSS windows inside the window; genes with no TSS inside their window (13) received no CAGE scores.

For each gene and track we also stored the map itself: the Gaussian-weighted mean of the unsigned and of the signed gradient (σ = 300 bp, ±1,200 bp) at the center of every 128-bp bin of the window (8,192 values). The cCRE scores were computed from the single-base gradients; spline interpolation of the stored values reproduced the exact scores of the CRISPRi-tested elements to within 0.1%.

#### Benchmark check

We computed the auPRC of each K562 and astrocyte track on the screens’ pairs whose gene is in the map (10,308 K562 pairs, 465 positives; 2,251 astrocyte pairs, 124 positives), scoring each tested element directly or by the highest-scoring registry cCRE overlapping it; pairs without a score took 0.

#### Enhancer–gene link cutoff

To call links we set one cutoff on the K562 screen [9]: the unsigned score at which the default K562 RNA-seq track (ENCODE total RNA-seq) recovered 70% (as for the ENCODE-rE2G map [9]) of the 465 CRISPRi positives of map genes, counting the 17 positives outside their gene’s window as missed. This cutoff, 2.02 × 10⁻³, was applied unchanged to every RNA-seq track; the equivalent for CAGE tracks was 1.23 × 10⁻³ (Supplementary Table 2). Precision and recall used the tested elements’ own scores, with 95% CIs from 10,000 bootstrap resamples of connected components of each screen’s element–gene graph. The astrocyte screen [10], not used to set the cutoff, served as a held-out test.

#### Biosample coverage

We compared E2GMap’s 454 biosamples (the distinct ontology terms of its 669 tracks) with the 369 terms of the 1,460 released ENCODE-rE2G prediction sets (ENCODE portal) and with the 252 cell types of the 640 primary cell conditions of the AlphaGenome 2 atlas (Supplementary Table 16 of Gayoso et al. [25]; each condition is a cell type within a tissue). Biosamples were grouped as cell types (Cell Ontology terms and other primary or in vitro differentiated cells), tissues (Uberon terms and other tissues), or cell lines. Two biosamples were counted as the same only if they had the same ontology term ID, after updating replaced IDs.

### Decomposing AlphaGenome’s attribution map via least squares

We fitted exact least squares to the log₂ unsigned scores of all cCRE–gene pairs and tracks, as a fixed-effects ANOVA with every two-way interaction between elements, genes, and tracks and no three-way interaction:

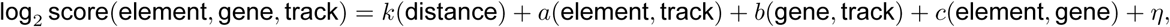

minimizing the sum of squared residuals *η* over all pairs and tracks. The distance is measured from the cCRE’s midpoint to the nearest TSS of the gene inside the window, across all of its GENCODE v50 transcripts (the window center for the 13 genes with none); *k* maps it to a reference 0–1 kb bin (*k* = 0) or to 29 log-spaced bins between 1 kb and 524 kb, with any longer distance in the last bin. The pair term *c* is constructed to average 0 within each distance bin, over each element’s genes, and over each gene’s elements. Altogether there are about 1.5 billion parameters, or about 10 observations per parameter.

Variance explained is the coefficient of determination (R²) of each track’s log₂ scores as the terms are added in the order distance, gene, element, pair; we report medians over the 669 tracks.

#### Contribution of each term to prediction

We used the K562 pairs whose gene is in the fit and whose tested element lies inside the gene’s window and overlaps at least one registry cCRE paired with the gene (6,593 pairs, 445 positives; the cCRE with the largest overlap stood for the element), and the astrocyte pairs selected in the same way (2,221 pairs, 124 positives). On these pairs we computed the auPRC of the distance term, then added the element, pair, and gene terms of the cell type’s RNA-seq or CAGE track in turn, and compared with the full score and with ABC. CIs are from 1,000 bootstrap resamples of 2-Mb blocks of genes, because nearby genes share cCREs and hence fitted terms.

#### Element term and measured chromatin

For the biosamples of the map’s tracks we collected ENCODE peak files for DNase-seq (204 biosamples), ATAC-seq (68), and H3K27ac ChIP-seq (152). A cCRE’s peak presence in a biosample was the fraction of the biosample’s files with an overlapping peak. For the default track of each biosample we subtracted from the element coefficients their mean over the default tracks of the assay, subtracted from peak presence its mean over the biosamples of the panel, and computed the Pearson correlation across cCREs with every biosample of the panel. We report the rank of the track’s own biosample (1, the highest correlation) and, as a null, its distribution over 1,000 permutations of biosample labels among tracks. A second DNase-seq panel added 15 biosamples that have no AlphaGenome 1 DNase-seq or ATAC-seq track, from 79 ENCODE files released on or after 19 September 2025.

#### Distance term and chromatin contact

We assigned to each cCRE–gene pair the balanced K562 intact Hi-C contact (ENCODE accession ENCFF621AIY; 5-kb bins) between the bins of the cCRE midpoint and of the gene’s nearest TSS, fitted log₂ contact of the 16.8 million pairs (76%) at least two 5-kb bins apart with non-zero contact, using the same 30 distance bins and element and gene terms, and compared the slopes of the distance terms (log₂ units per doubling of distance) over the 17 bins between 10.8 and 422 kb.

### Reproducibility

We ran our benchmarking experiments on 2 × NVIDIA RTX 6000 Ada Generation GPUs and an NVIDIA Quadro RTX 4000. The genome-wide enhancer–gene mapping was done on NVIDIA H200 GPUs.

For Enformer, we used the community PyTorch port from https://github.com/lucidrains/enformer-pytorch (v0.8.10) with the EleutherAI/enformer-official-rough checkpoint. For Borzoi, we used the community PyTorch port from https://github.com/johahi/borzoi-pytorch (v0.5.1) with the four replicate checkpoints (johahi/borzoi-replicate-0..3). For AlphaGenome, we used the community PyTorch port from https://github.com/genomicsxai/alphagenome-pytorch (commit 473d3d2) with weights locally converted from the publicly released alphagenome-jax all_folds-v1 checkpoint (670/670 parameter tensors). For Flashzoi, we used johahi/borzoi-pytorch (flashed=True) with the four replicate checkpoints johahi/flashzoi-replicate-0..3 at Hugging Face revisions 72040fb, 3e54f34, 524b05b, and 336e45f; Flashzoi’s authors run the model in mixed precision, and we re-implemented its attention in fp32. For Cerberus, we used Calico’s code from calico/baskerville (releases/cerberus, commit 34bfe56) and its eight fold models (f0c0–f7c0, from gs://seqnn-share/cerberus/models), with PyTorch 2.11.0, CUDA 12.8, Triton 3.6.0, and mamba_ssm 2.3.2.post1, run with TRITON_F32_DEFAULT=ieee. For NTv3, we used InstaDeep’s PyTorch modeling code (InstaDeepAI/ NTv3_pretrained, revision 6966543) and the access-gated NTv3_tracks_1mb weights (revision 9188406), both checksum-verified, with PyTorch 2.13.0 and CUDA 12.6. TF32 was disabled for Enformer, Borzoi, Flashzoi, Cerberus, and NTv3.

For ENCODE-rE2G, we refitted the logistic regression models with the settings of ENCODE_rE2G v1.0.0 (https://github.com/EngreitzLab/ENCODE_rE2G) on the K562 feature tables of https://github.com/karbalayghareh/ENCODE-E2G (commit 7b4a653): the DNase-only table for rE2G-DNase and the all-features table for rE2G-extended, which lacks numCandidateEnhGene, so rE2G-extended was refitted on the other 44 of its 45 features. For astrocytes, the rE2G-DNase features were taken from the ENCODE-rE2G astrocyte predictions (ENCFF440FMQ). For EGrf, we used the K562 and astrocyte feature tables of Green et al. [10] from https://github.com/Voineagulab/astrocyte_crispri (commit ec6e4e0) and refitted the random forests (1,000 trees) with scikit-learn 1.6.1.

The K562 CRISPRi screen [9] was obtained from https://github.com/EngreitzLab/CRISPR_comparison/blob/ v1.0.0/resources/crispr_data/EPCrisprBenchmark_combined_data.training_K562.GRCh38.tsv.gz. The NHA CRISPRi screen [10] was obtained from https://github.com/Voineagulab/astrocyte_crispri (commit ec6e4e0; the 2,307-pair modeling table and the full results for 7,759 pairs). ABC scores in K562 are those released with the ENCODE-rE2G feature table [9], and ABC scores in astrocytes are those of the ENCODE astrocyte predictions (ENCFF440FMQ), which use DNase-seq and Hi-C averaged over cell types; pairs without a score were set to 0.

## Code and data availability

Our enhancer–gene scoring framework E2GScore, which includes the genome-wide mapping code, is available at https://github.com/cthe4801/e2gscore. It contains the configuration files for every reported experiment, the enhancer–gene scores behind the reported numbers, the five CRISPRi benchmark tables, the FlowFISH+ and Gasperini pair tables and gene annotation provided by the authors of Linder et al. [23], and a ledger mapping each result table to its experimental configuration.

E2GScore can be run for a model implemented in any machine learning framework and on different machines by modifying the backend_<model<.py file, which requires the model’s forward pass (and backward pass for input gradients) to be implemented, while the scoring algorithm in score_eg.py does not change.

Our genome-wide enhancer–gene map is available at https://e2gscore.com. AlphaGenome-derived scores and figures (processed as described in Methods), including E2GMap, are subject to the AlphaGenome Output Terms of Use (https://deepmind.google.com/science/alphagenome/output-terms), not this article’s CC BY license.

## Acknowledgements

This research was supported by the Commonwealth through an Australian Government Research Training Program (RTP) Scholarship (DOI: https://doi.org/10.82133/C42F-K220) and by a University Postgraduate Award (UPA) from the University of New South Wales (UNSW Sydney) to C.T. The research was also supported by an NHMRC Investigator Grant to I.V. This research includes computations using the HPC cluster Katana supported by Research Technology Services at UNSW Sydney (https://doi.org/10.26190/669x-a286).

## Competing interests

The authors declare no competing interests.

## Use of large language models

We used large language models (Claude Opus 5.5, Claude Fable 5.2, ChatGPT Sol 5.6) for data preprocessing, coding, and analysis, as well as manuscript drafting, formatting, and review. The authors take full responsibility for all results.

## Supplementary Figures

**Supplementary Fig. 1.**
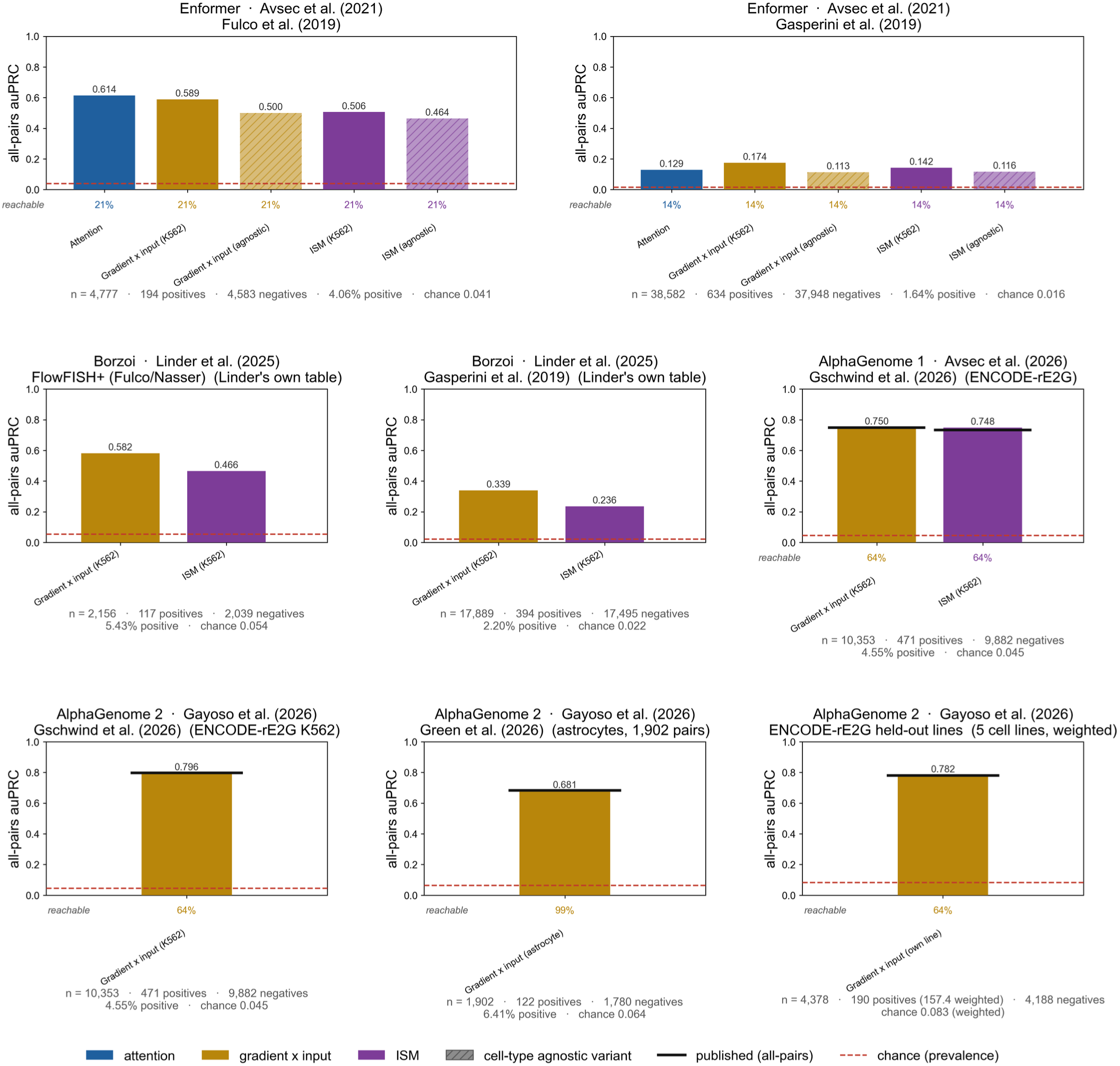
All-pairs auPRC of reproduced published benchmarking results for Enformer, Borzoi, and AlphaGenome 1 and 2. Top, Enformer [22] on the Fulco (4,777 pairs, 194 positives) and Gasperini (38,582; 634) cohorts. Middle, Borzoi [23] on the authors’ FlowFISH+ (2,156; 117) and Gasperini (17,889; 394) tables (all pairs inside Borzoi’s input window), and AlphaGenome 1 [24] on the ENCODE-rE2G K562 benchmark (10,353; 471). Bottom, AlphaGenome 2 [25] on that benchmark, the astrocyte evaluation subset (1,902; 122), and the ENCODE-rE2G held-out benchmark (4,378; 190; weighted auPRC, as in Supplementary Fig. 4e). Bars: attention (blue), gradient × input (gold), ISM (purple); hatched, cell-type-agnostic readout; black lines, published all-pairs values (none for Enformer and Borzoi); dashed lines, prevalence; %, share of pairs inside the input window. Enformer bars are computed without the ±57,344-bp crop, genomic replacement, and 16-token rule of Supplementary Fig. 2.

**Supplementary Fig. 2.**
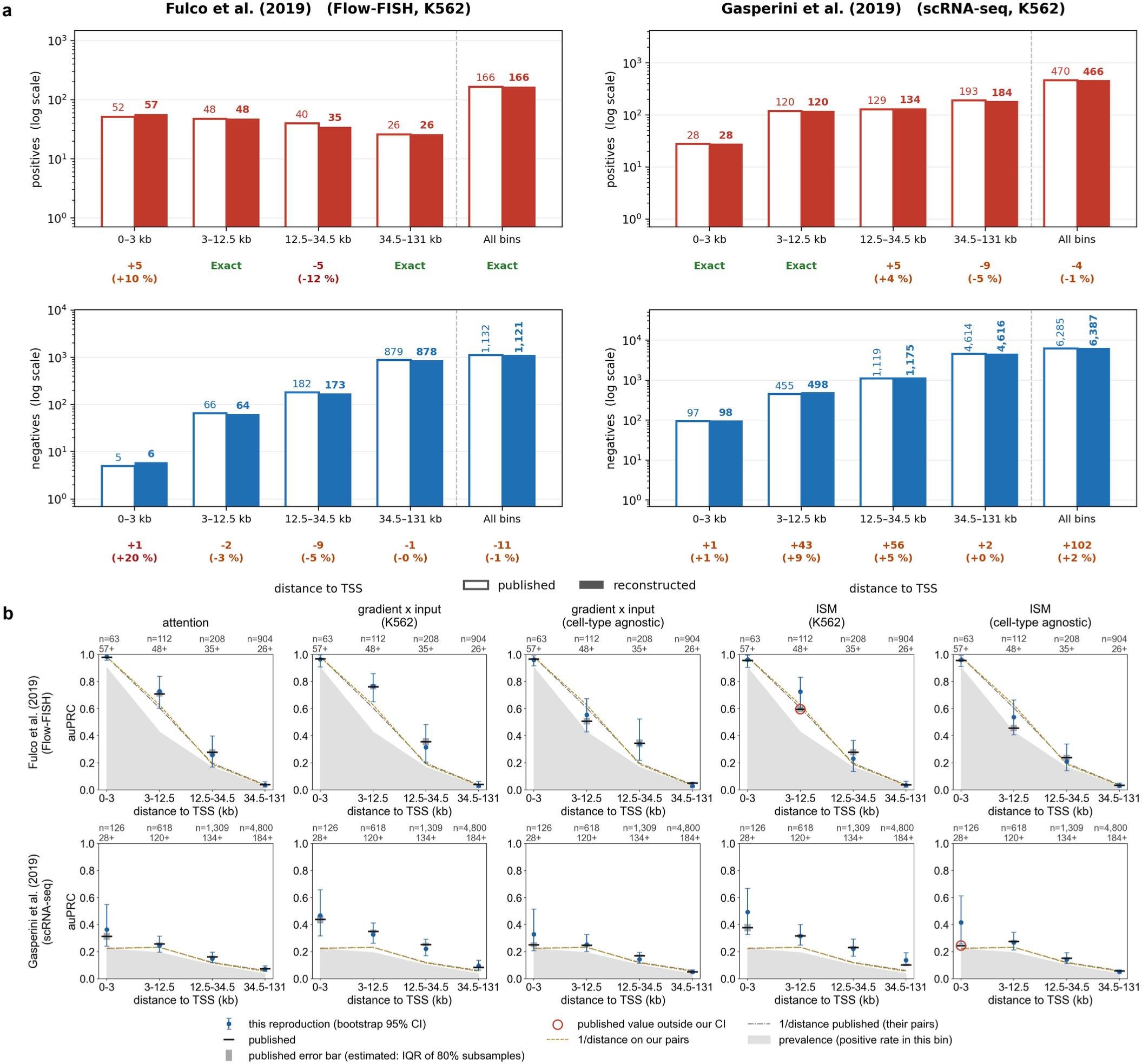
E2GScore reproduces 38 of the 40 published Enformer auPRCs of Avsec et al. [22]. **a**, Positive (top) and negative (bottom) pairs per distance bin in the rebuilt Fulco (left) and Gasperini (right) cohorts (filled) and as published (open); counts agree to 98.2% (1,287 rebuilt against 1,298 published pairs, 23 misplaced; 166 positives in both) and 98.3% (6,853 against 6,755, 116 misplaced; 466 against 470 positives). **b**, auPRC per distance bin for attention, gradient × input, and ISM with K562 or cell-type-agnostic readouts (their Fig. 2b and Extended Data Fig. 7c). Points, this reproduction with 95% bootstrap intervals over pairs (10,000 resamples per bin); black bars, published values; gray bars, published error bars (interquartile range over 80% subsamples, estimated); red circles, published values outside our intervals; dashed and dash-dot lines, 1/distance auPRC on our pairs and as published; gray fill, prevalence; *n* and +, pairs and positives.

**Supplementary Fig. 3.**
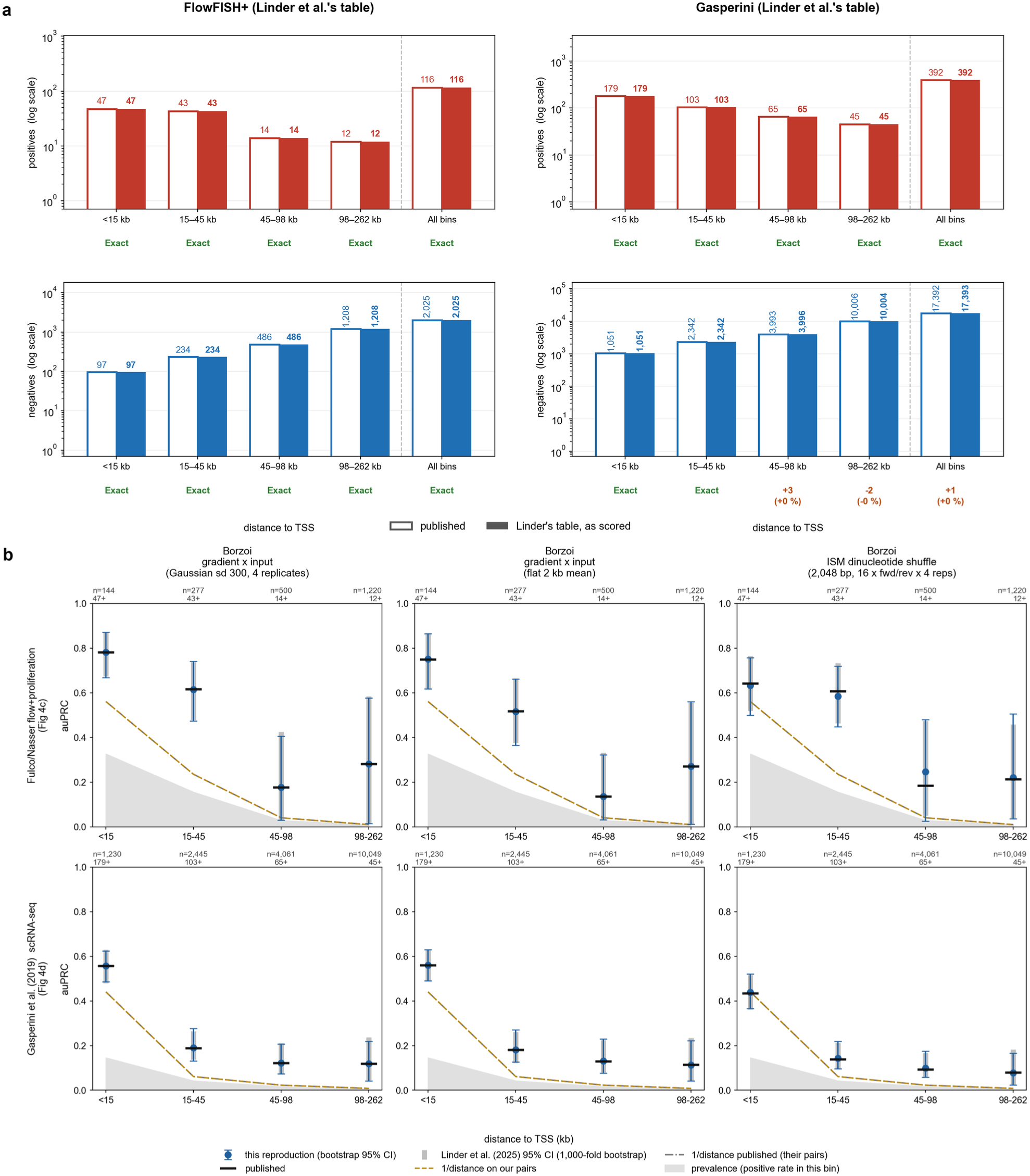
E2GScore reproduces all 24 published Borzoi auPRCs of Linder et al. [23]. **a**, Positive (top) and negative (bottom) pairs per distance bin in the FlowFISH+ (left; 2,141 pairs) and Gasperini (right; 17,785) tables, counting the pairs inside the four distance bins, as scored (filled) and as published (open; 2,141 and 17,784; 0 and 5 pairs misplaced; 116 and 392 positives in both). **b**, auPRC per distance bin, four replicates: gradient × input under a Gaussian (σ = 300 bp; their Fig. 4c,d) or averaged over a flat 2-kb window, and ISM (2,048 bp shuffled as dinucleotides 16 times per orientation and replicate; signed mean log change; their Supplementary Fig. 7b). Points, this reproduction with 95% bootstrap intervals over pairs (1,000 resamples per bin as drawn; the counts in the text use 10,000 resamples, Supplementary Table 2); black bars, published values; gray bars, published 95% intervals; dashed line, 1/distance auPRC (ours and the published coincide); gray fill, prevalence; *n* and +, pairs and positives.

**Supplementary Fig. 4.**
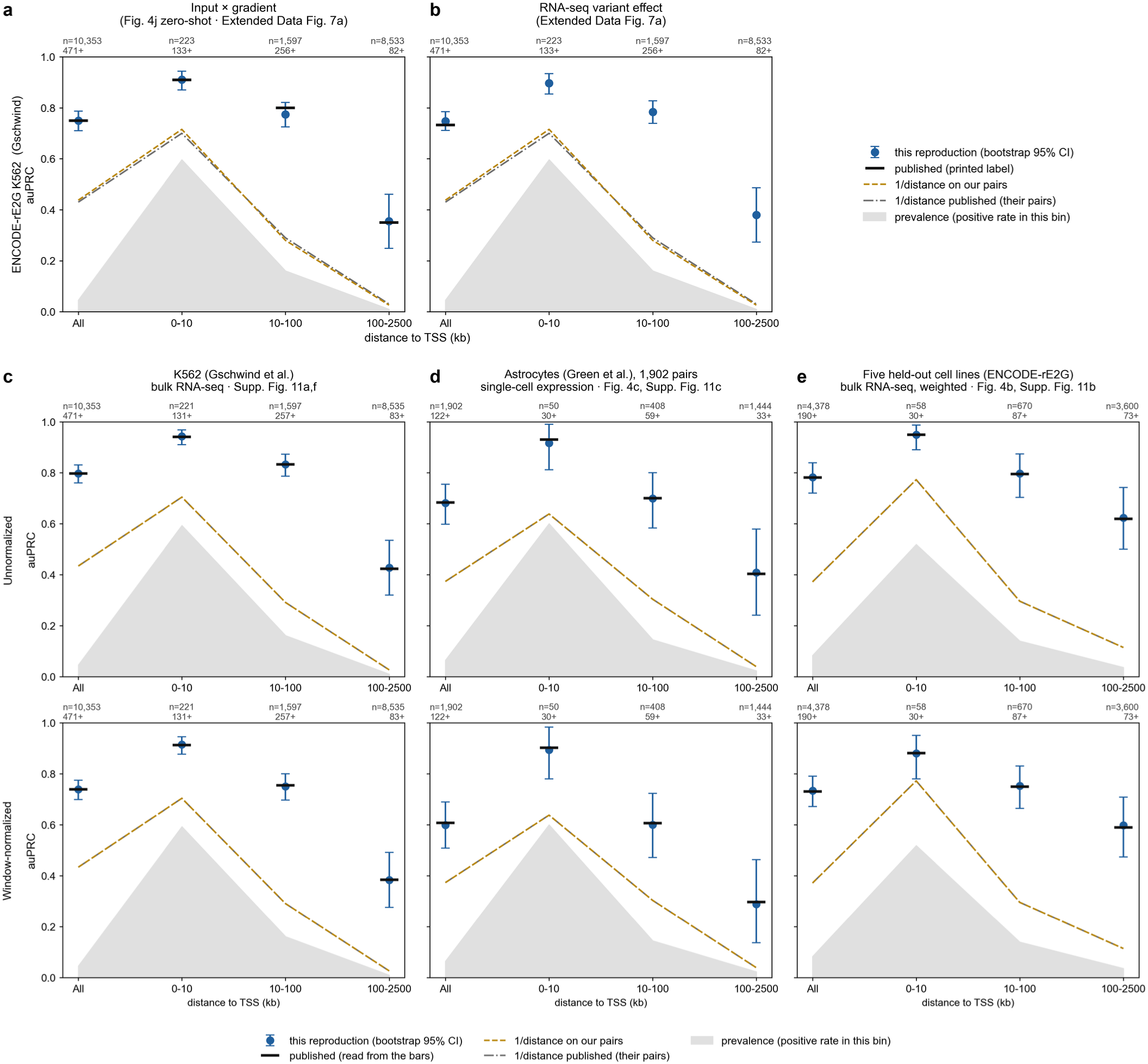
E2GScore reproduces all five published auPRCs of AlphaGenome 1 and all 24 of AlphaGenome 2. auPRC for all pairs and three distance bins (each paper’s TSS distances); panel titles cite the reproduced papers’ figures. **a**,**b**, AlphaGenome 1 [24] scored with E2GScore on the ENCODE-rE2G K562 benchmark [9] by input × gradient (**a**) or RNA-seq variant effect, a whole-element ISM (**b**; only its all-pairs value is published). **c**–**e**, AlphaGenome 2 [25] input gradients, unnormalized (top) or window-normalized (bottom), scored with the AlphaGenome API client’s scoring function on the same K562 benchmark (**c**), the astrocyte evaluation subset (**d**), and the ENCODE-rE2G held-out benchmark (**e**; auPRC weighted by each pair’s direct-effect probability). Points, this reproduction with 95% bootstrap intervals over pairs (2,000 resamples per bin in **a**,**b**; 10,000 in **c**–**e** and in Supplementary Table 2); black bars, published values (read from the bars in **c**–**e**); dashed and dash-dot lines, 1/distance auPRC, ours and as published (coincident in **c**–**e**); gray fill, prevalence; *n* and +, pairs and positives per bin.

**Supplementary Fig. 5.**
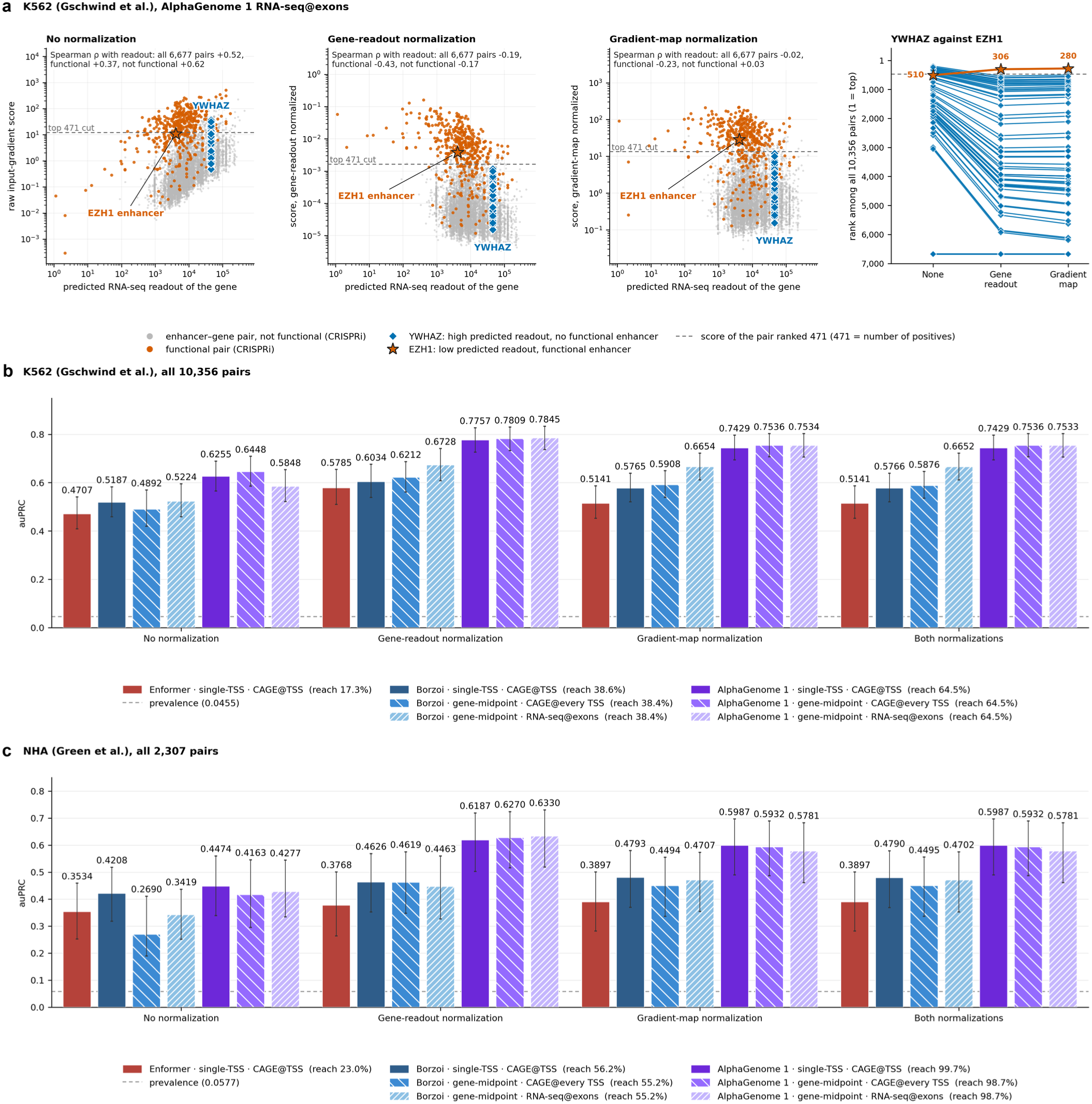
Normalizing input gradient scores across genes raises auPRC in both screens. **a**, K562, AlphaGenome 1 RNA-seq@exons (6,677 in-window pairs): raw, gene-readout-normalized, and gradient-map-normalized scores against the gene’s predicted readout (orange, functional pairs; gray, non-functional; dashed line, score of the pair ranked 471, the number of positives; ρ, Spearman correlation), and the rank among all 10,356 pairs of the tested pairs of *YWHAZ* (no functional enhancer; diamonds) and of a functional *EZH1* enhancer (star). **b**,**c**, auPRC over all pairs in K562 (**b**; 10,356 pairs, 471 positives) and NHA (**c**; 2,307; 133) for seven readouts of Enformer, Borzoi, and AlphaGenome 1 under no normalization, gene-readout normalization, gradient-map normalization, or both (equivalent to gradient-map normalization; Methods); pairs outside the input window score 0. Error bars, 95% CI (connected-component bootstrap, 10,000 resamples); reach, share of pairs inside the input window; dashed line, prevalence.

**Supplementary Fig. 6.**
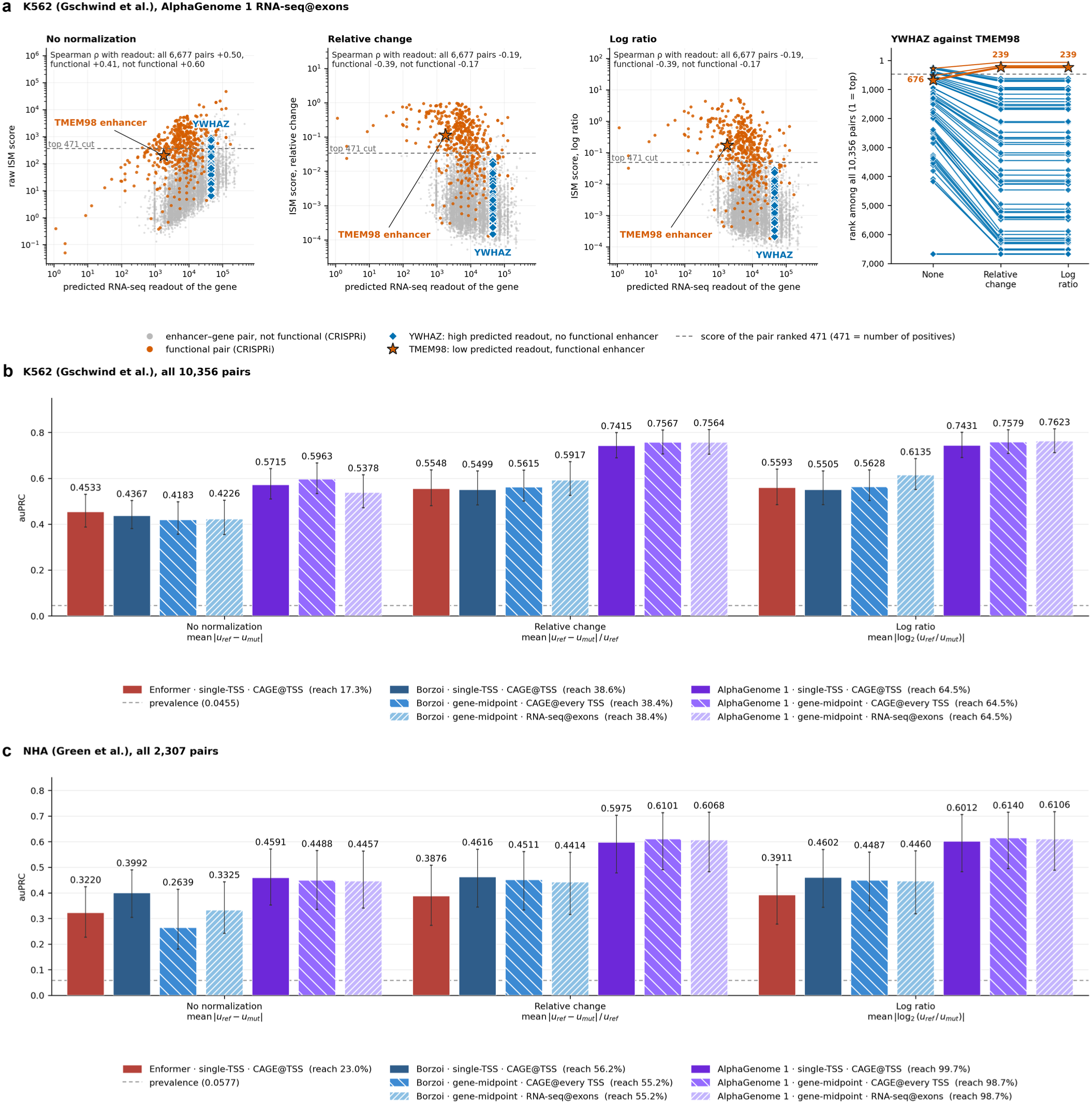
Normalizing ISM scores across genes raises auPRC in both screens. **a**, As Supplementary Fig. 5a, for ISM (here *u* is the summed readout *S* of equation (1)): the raw score (mean |*u*_ref_ − *u*_mut_| over ten shuffles), the relative change (mean |*u*_ref_ − *u*_mut_| / *u*_ref_), and the log ratio (mean |log_2_(*u*_ref_ / *u*_mut_)|) against the gene’s predicted readout *u*_ref_, and the rank among all 10,356 pairs of the tested pairs of *YWHAZ* (diamonds) and of the three functional enhancers of *TMEM98* (stars; large, the strongest CRISPRi effect). **b**,**c**, auPRC over all pairs in K562 (**b**) and NHA (**c**) for the seven readouts of Supplementary Fig. 5 under the three statistics of **a**: no normalization, relative change, and log ratio (the score used in Supplementary Fig. 10). Error bars, 95% CI (connected-component bootstrap, 10,000 resamples); reach, share of pairs inside the input window; dashed line, prevalence.

**Supplementary Fig. 7.**
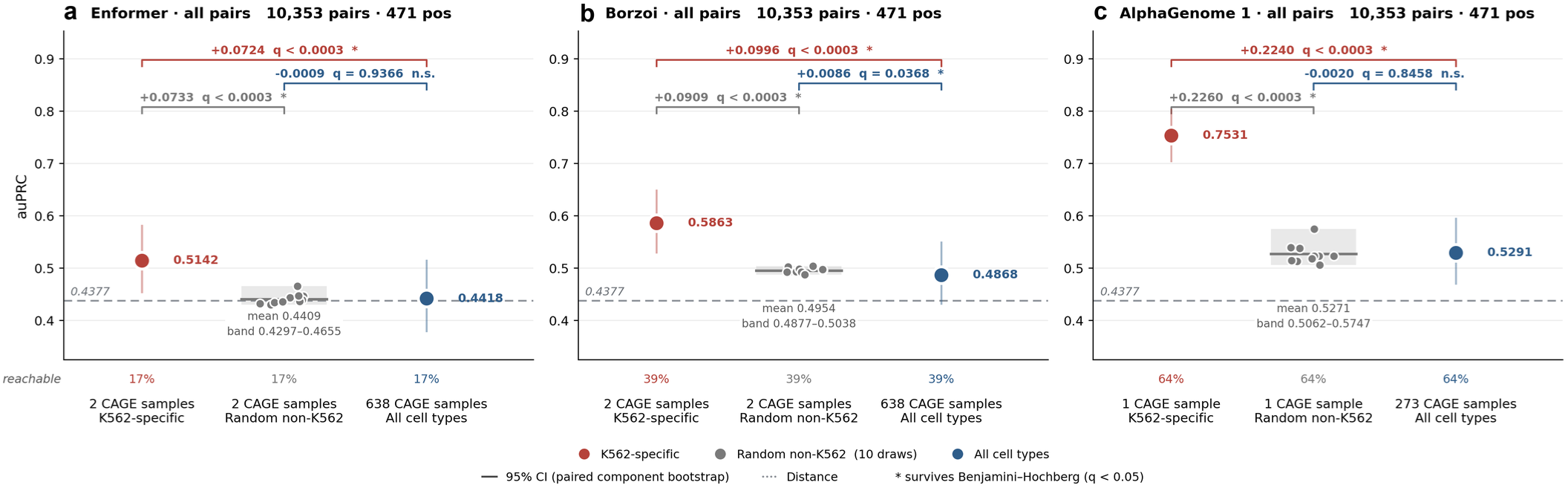
Cell-type-matched CAGE tracks improve enhancer–gene linking in K562. auPRC of input gradients for Enformer (**a**), Borzoi (**b**), and AlphaGenome 1 (**c**), with the CAGE readout taken from the K562 tracks (red), from ten random draws of as many non-K562 tracks (gray points; line, mean; shading, range), or from all CAGE tracks (blue), on all 10,353 K562 pairs [9], with pairs outside a model’s input window scored 0 (panel headings: pos, positives). Reachable, share of the pairs inside the model’s input window (these hold 389, 432, and 454 of the 471 positives). Dashed line, auPRC of 1/distance on the same pairs. Error bars, 95% CI (connected-component bootstrap, 10,000 resamples); brackets, paired differences (left minus right; random draws, their mean) with Benjamini–Hochberg q over the nine comparisons shown. The input window is centered on, and CAGE is read over the 384 bp around, the RefSeq TSS given in the K562 table instead of the MANE Select TSS (Supplementary Fig. 8); scores use gradient-map instead of gene-readout normalization (Supplementary Fig. 5).

**Supplementary Fig. 8.**
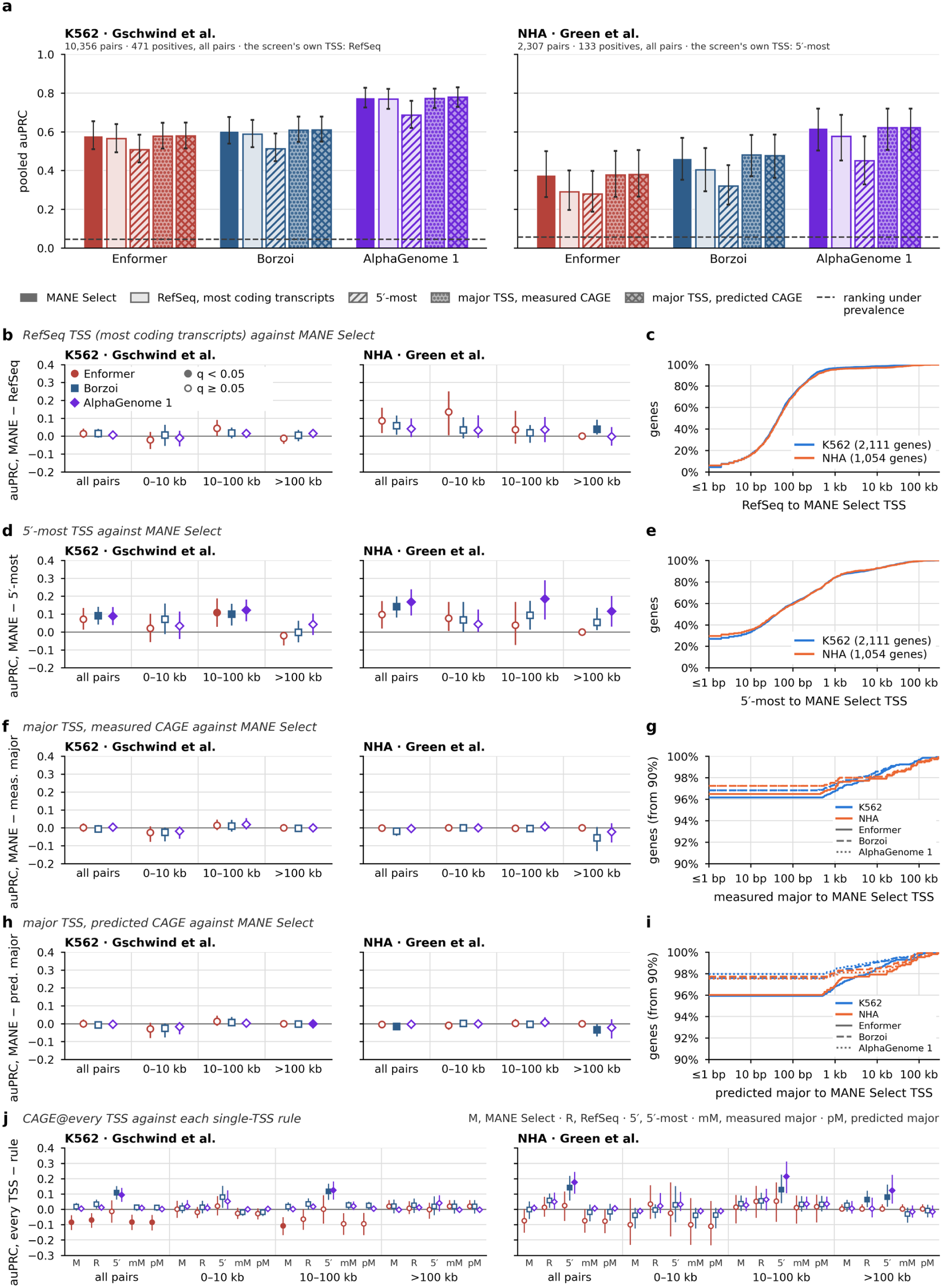
CAGE at the MANE Select TSS matches the major-TSS selection rules and outperforms the 5′-most TSS for Borzoi and AlphaGenome 1. **a**, Pooled auPRC of Enformer, Borzoi, and AlphaGenome 1 over all pairs (K562, 10,356 pairs, 471 positives; NHA, 2,307; 133) with CAGE read at, and the window centered on, one TSS: MANE Select; the RefSeq TSS shared by the most coding transcripts (the K562 table’s TSS); the 5′-most GENCODE v32 TSS (the NHA table’s TSS); or the major TSS, the GENCODE v50 TSS with the highest measured (FANTOM5) or predicted CAGE signal in the screen’s cell type. Dashed lines, random ranking under prevalence. **b**,**d**,**f**,**h**, auPRC with MANE Select minus RefSeq (**b**), 5′-most (**d**), measured major (**f**), or predicted major (**h**), over all pairs and by distance; filled, Benjamini–Hochberg q < 0.05 (Supplementary Table 2). **c**,**e**,**g**,**i**, Cumulative share of genes (K562, 2,111; NHA, 1,054 in **c**,**e** and 1,053 in **g**,**i**) by distance from that TSS to the MANE Select TSS. **j**, auPRC with CAGE@every TSS minus CAGE at each single TSS of **a** (M, MANE Select; R, RefSeq; 5′, 5′-most; mM, measured major; pM, predicted major), over all pairs and by distance; filled, Benjamini–Hochberg q < 0.05 (Supplementary Table 2; column M takes the q of Fig. 2). CAGE@every TSS, with the window centered on the gene midpoint, is the readout of Fig. 2. Error bars, 95% CI (connected-component bootstrap, 10,000 resamples).

**Supplementary Fig. 9.**
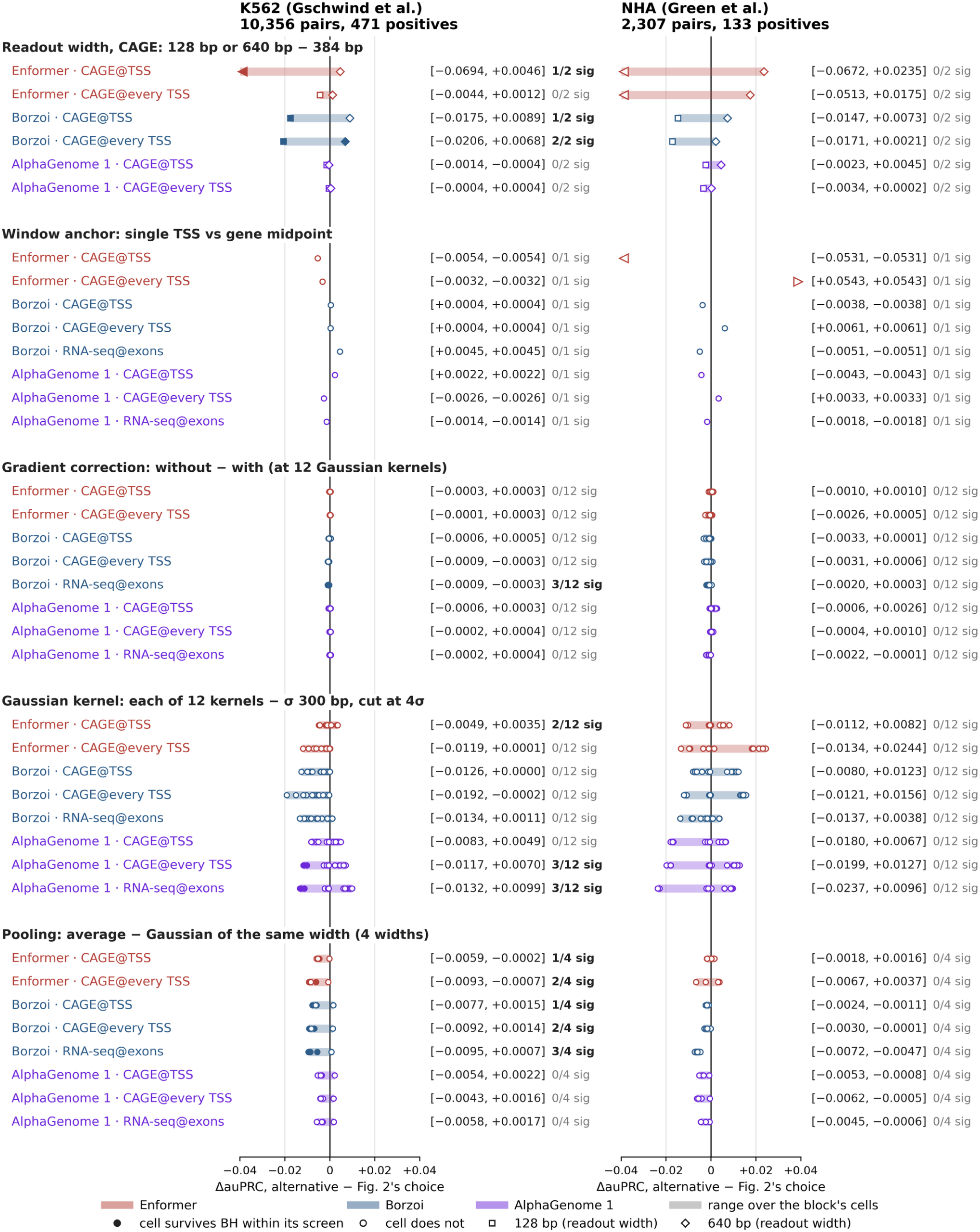
Varying other protocol parameters has minor to no effects on model performance. Difference in auPRC, alternative minus the choice of Fig. 2, for the eight readouts of Enformer, Borzoi, and AlphaGenome 1 in K562 (left) and NHA (right), on each readout’s reachable pairs, paired on Fig. 2’s 10,000 bootstrap resamples. Readout width (the six CAGE readouts): CAGE read over 128 bp (squares) or 640 bp (diamonds) at the MANE Select TSS (CAGE@TSS) or at every annotated TSS (CAGE@every TSS), minus Fig. 2’s 384 bp. Window anchor: gene-midpoint minus single-TSS window (CAGE@TSS), or single-TSS minus gene-midpoint window (CAGE@every TSS and RNA-seq@exons), on pairs inside both windows. Gradient correction: uncorrected minus corrected, at each of 12 Gaussian kernels (matched in width to flat windows of 500, 1,000, 2,000, or 3,000 bp; cut at 1σ, 2σ, or 3σ). Gaussian kernel: each of these minus the protocol kernel (σ = 300 bp, cut at 4σ). Pooling: flat-window average minus the Gaussian of the same width. Each marker is one comparison (square, 128-bp readout; diamond, 640-bp readout; dot, all others). Filled markers are significant (Benjamini–Hochberg q < 0.05 over each screen’s 244 comparisons). The shaded bar and the bracketed numbers give the range of a row’s comparisons, and “k/N sig” the number of them that are significant. A triangle marks a value beyond the axis.

**Supplementary Fig. 10.**
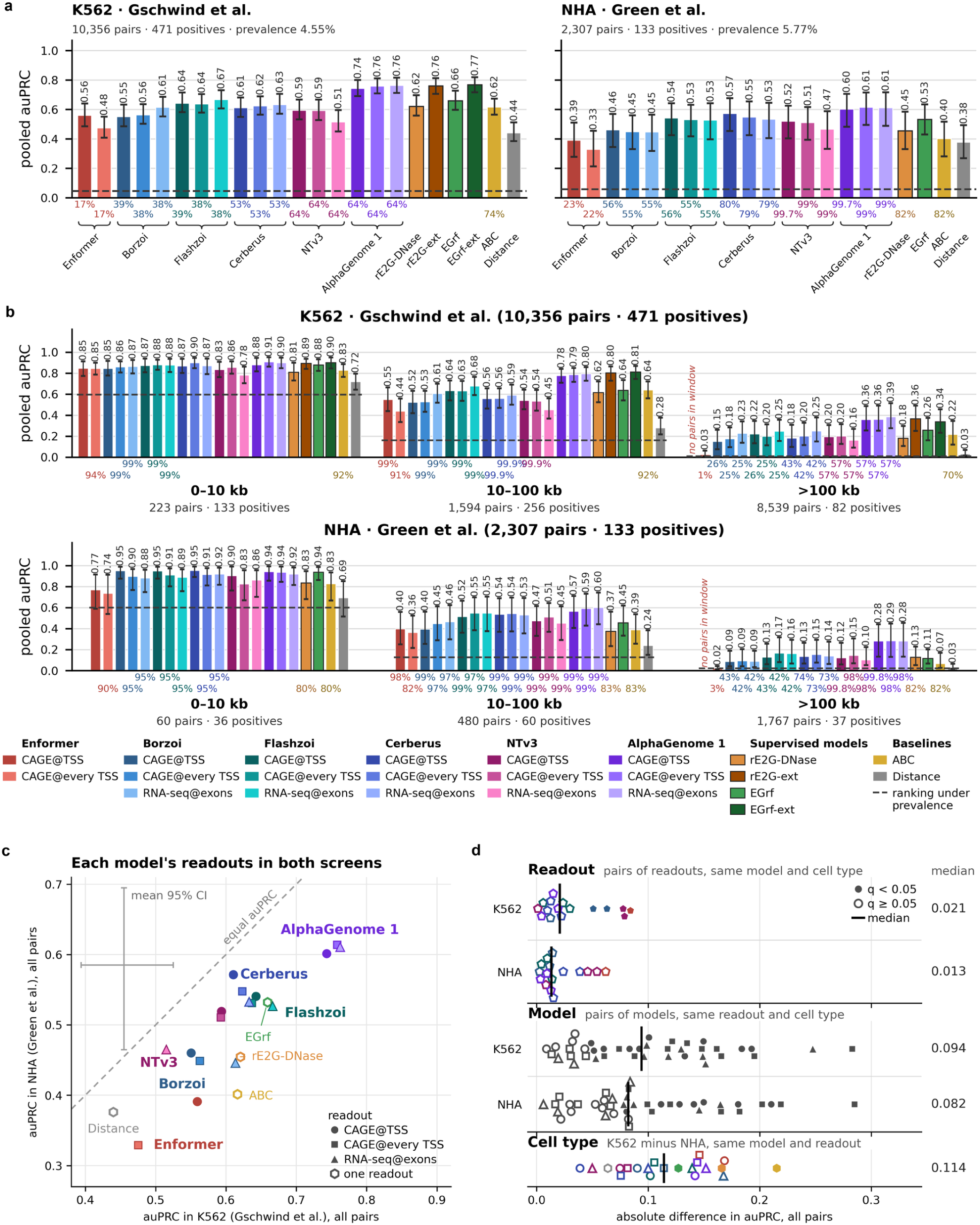
Under matched protocols, model rankings for perturbation-based attribution scores are consistent across gene readouts and cell types. **a**–**d**, As Fig. 2a–d (same pairs, bootstrap resamples, colors, and symbols), with every seq2func model scored by ISM instead of input gradients: the mean absolute change in the log₂ gene readout when the 2,000 bp centered on the enhancer are replaced by each of ten dinucleotide-preserving shuffles [51]. AlphaGenome 2 is not shown; the expert models and Distance are those of Fig. 2. Over all pairs, ISM minus input gradients was −0.089 to −0.019 auPRC for the 17 K562 readouts (*q* < 0.05 for 15) and −0.044 to +0.028 for the 17 NHA readouts (q > 0.26; Benjamini–Hochberg over each screen’s 292 comparisons). **d**, *n* = 16 comparisons between readouts and 40 between models per screen, and 21 between the screens.

**Supplementary Fig. 11.**
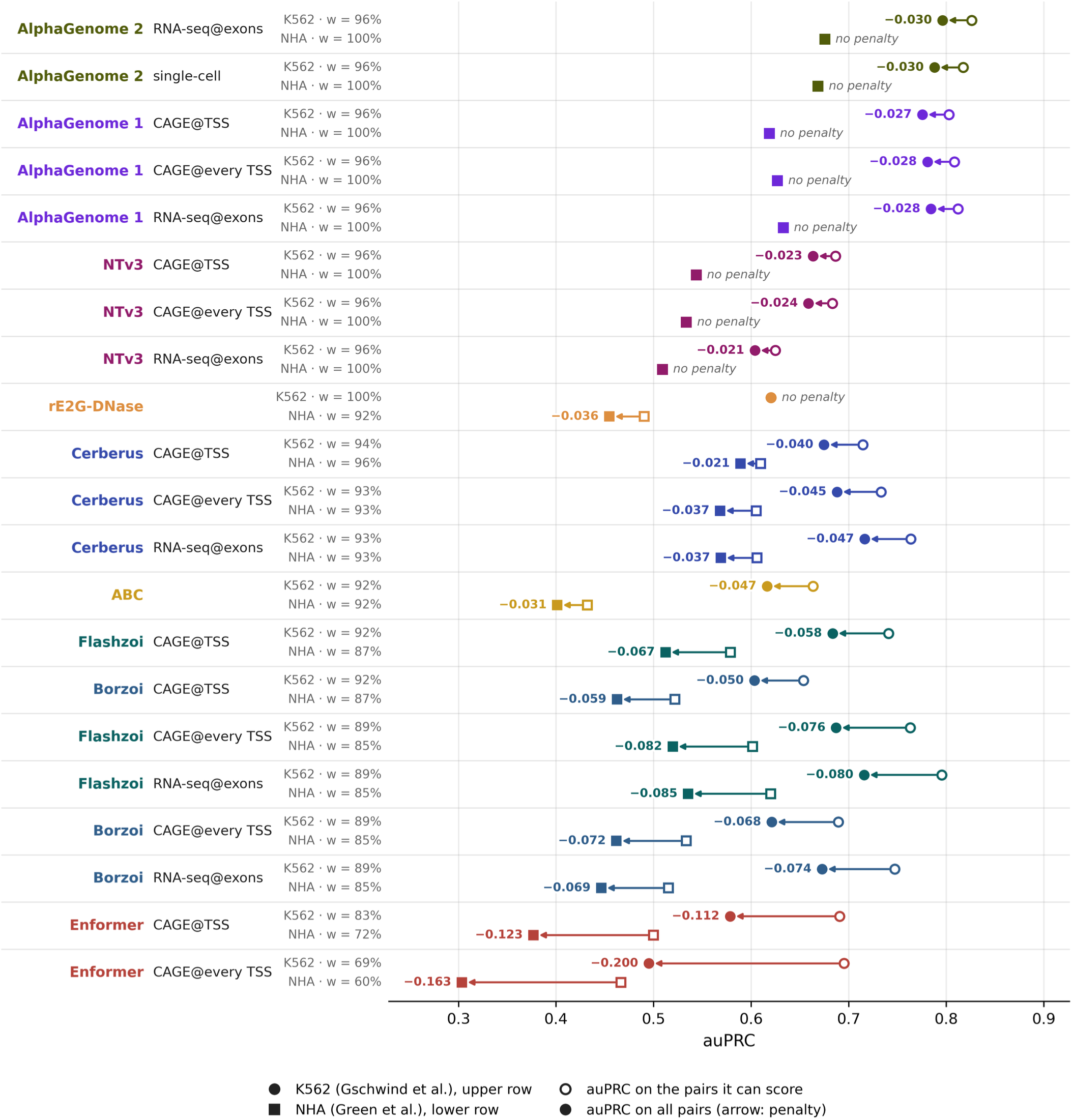
Methods that score fewer positives lose more auPRC on the pairs they cannot score. For each method in K562 (circles, upper row; prevalence 4.5%) and NHA (squares, lower row; prevalence 5.8%): open marker, auPRC on the pairs the method can score; filled marker, auPRC on all pairs, where the pairs it cannot score tie at the bottom; *w*, share of positives it can score. Under tie-aware average precision, auPRC on all pairs = w × auPRC on scorable pairs + (1 − w) × prevalence, so the arrow, the reach penalty, equals (1 − w) × (auPRC on scorable pairs − prevalence). Methods are sorted by w (mean of the two screens), highest first. rE2G-ext, EGrf, EGrf-ext, and Distance score every pair and are not shown. Scorable pairs differ between methods, so open markers are not comparable across methods.

**Supplementary Fig. 12.**
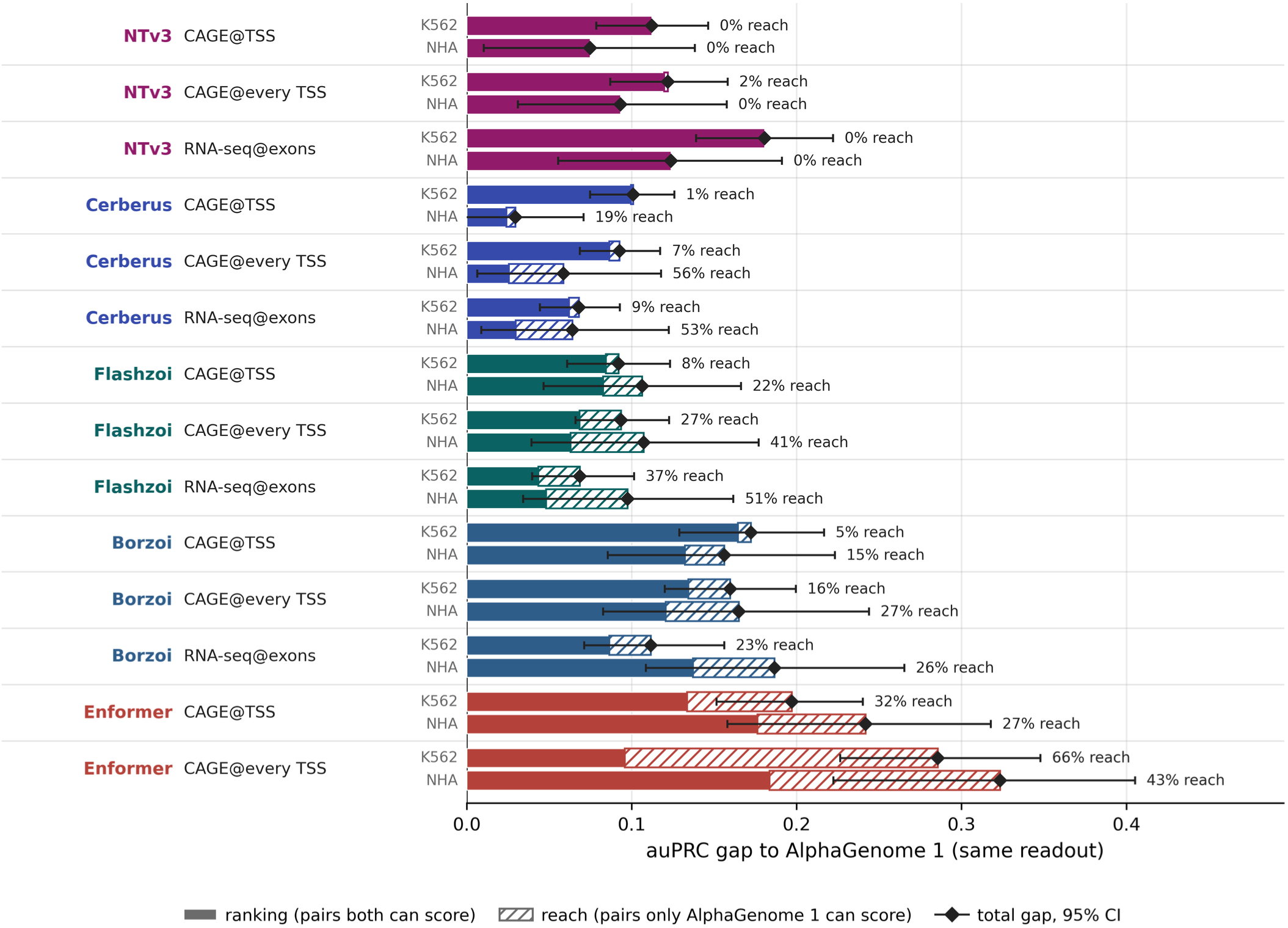
Most of AlphaGenome 1’s advantage over the other seq2func models comes from ranking the pairs both can score. For each Enformer, Borzoi, Flashzoi, Cerberus, and NTv3 readout, the auPRC gap to AlphaGenome 1 with the same readout in K562 (upper row) and NHA (lower row) is split exactly into ranking (filled bar: the auPRC difference on the pairs both models can score, weighted by their share of positives) and reach (hatched bar: the contribution of the pairs only AlphaGenome 1 can score). Labels, reach as a percentage of the gap. Diamonds and whiskers, the gap and its 95% CI (10,000 bootstrap resamples of connected components, as in Fig. 2). Models are ordered by *w*, the share of positives they can score, highest first. Reach exceeds half of the gap in 4 of the 28 comparisons. For Cerberus CAGE@TSS in NHA, the 95% CI of the gap (−0.015 to 0.071) extends below the axis.

**Supplementary Fig. 13.**
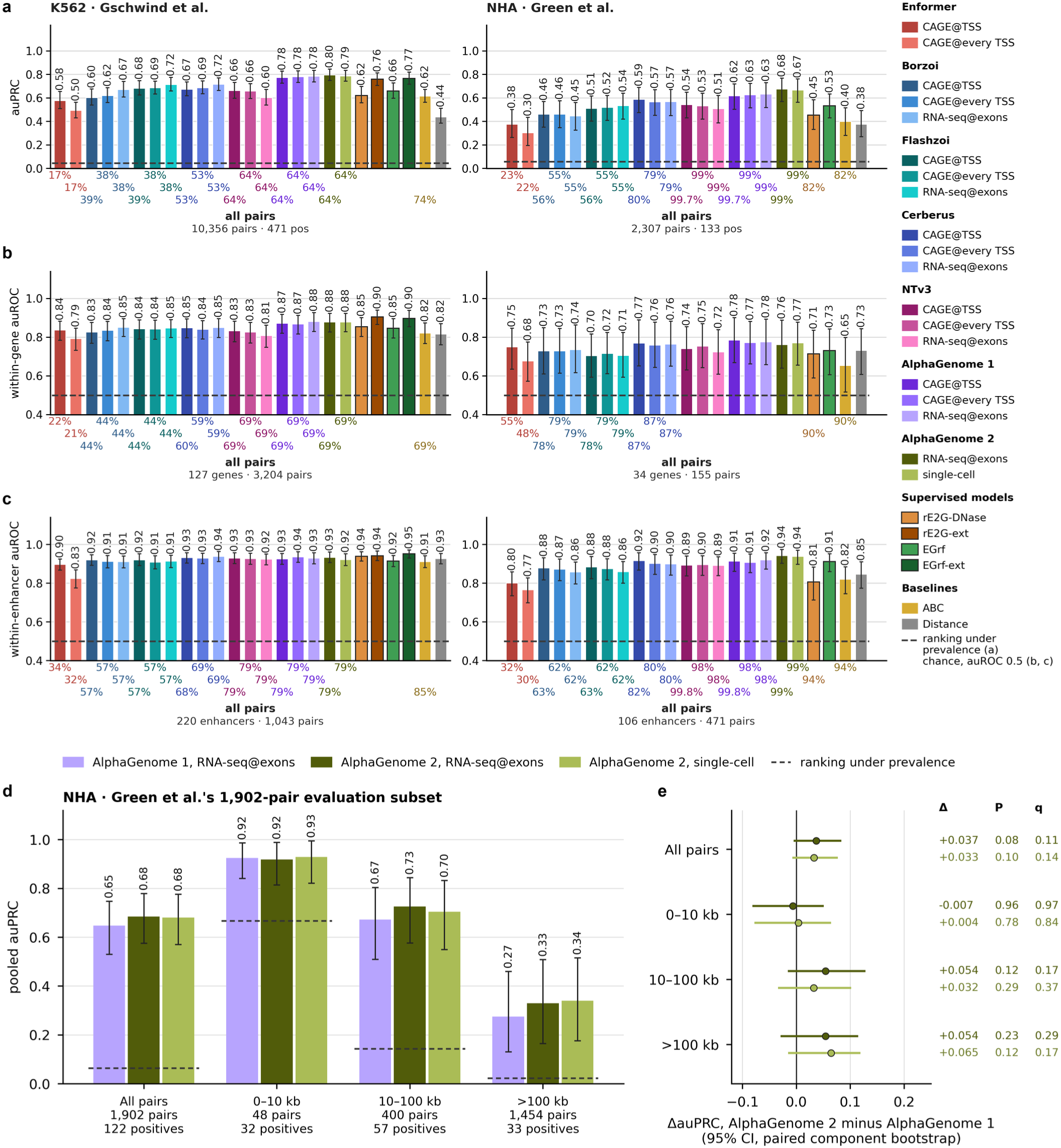
Pooled, within-gene, and within-enhancer rankings of every method over all pairs, and AlphaGenome 2 against AlphaGenome 1 on another astrocyte evaluation subset. **a**–**c**, Every method of Fig. 2 on all pairs of the K562 (left) [9] and NHA (right) [10] screens. **a**, Pooled auPRC (as in Fig. 2a); the pairs and positives (pos) are given below each panel. **b**, Macro within-gene auROC (area under the receiver operating characteristic curve): the auROC of each gene’s pairs, averaged over the genes with at least one positive and one negative pair. **c**, Macro within-enhancer auROC: the auROC of each enhancer’s pairs, averaged over the enhancers with at least one positive and one negative pair. In **b** and **c**, the numbers of such genes or enhancers and of their pairs are given below each panel; NHA has only 34 such genes (K562, 127), so its intervals in **b** are wide. Error bars, 95% CI (10,000 bootstrap resamples of connected components). Percentages, share of the pairs a method can score (in **b** and **c**, of the pairs that enter the average), shown when below 100%. Dashed lines, random ranking under prevalence (**a**) and chance, auROC 0.5 (**b**, **c**). No significance tests were performed for **b** and **c**. **d**, Pooled auPRC on the 1,902-pair evaluation subset of Green et al. [10] (122 positives) evaluated by Gayoso et al. [25], over all pairs and by distance to the MANE Select TSS, for AlphaGenome 1 (RNA-seq@exons) and for AlphaGenome 2 scored through its API (bulk RNA-seq over exons, and single-cell expression). Dashed lines, random ranking under prevalence. **e**, AlphaGenome 2 minus AlphaGenome 1 with its 95% CI, two-sided bootstrap P, and Benjamini–Hochberg q over the 144 comparisons of AlphaGenome 2’s two readouts with the 18 other readouts and methods on this subset, over all pairs and in three distance bins. Error bars and CIs in **d** and **e**, 10,000 bootstrap resamples of the subset’s 278 connected components.

**Supplementary Fig. 14.**
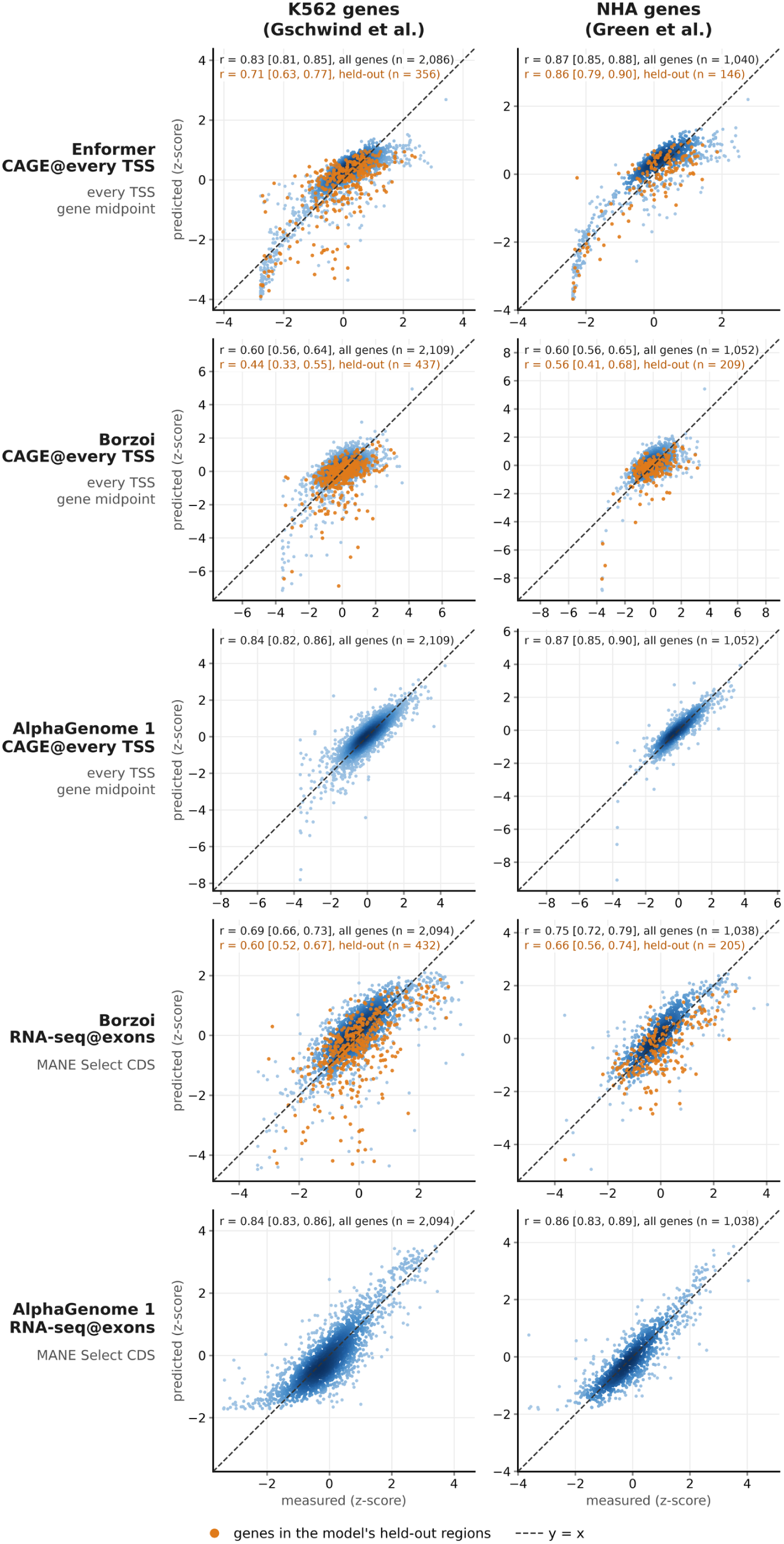
Enformer, Borzoi, and AlphaGenome 1 predict the expression of the screens’ genes at least as accurately in astrocytes as in K562. Predicted against measured expression of the genes tested in K562 (left) [9] and astrocytes (NHA, right) [10]. One point per gene; axes, log₂(1 + x), z-scored within each panel; dashed line, y = x. Rows 1–3, CAGE@every TSS (as in Fig. 2) against FANTOM5 CAGE in the same bins. Rows 4–5, RNA-seq averaged over the MANE Select coding sequence (MANE Select CDS) against bulk RNA-seq TPM (K562, ENCODE ENCSR000AEM; NHA, Green et al. [10]); protein-coding genes only (2,094 of the 2,109 K562 genes and 1,038 of the 1,052 astrocyte genes). *r*, Pearson correlation with 95% confidence interval (2,000 gene-resampling bootstrap draws); *n*, genes (Enformer cannot read genes lacking a TSS in its predicted 114,688 bp). Orange, genes (and r) in the model’s held-out regions (AlphaGenome 1 was trained on all folds). The CAGE libraries were among the models’ training data; no model saw the NHA RNA-seq.

**Supplementary Fig. 15.**
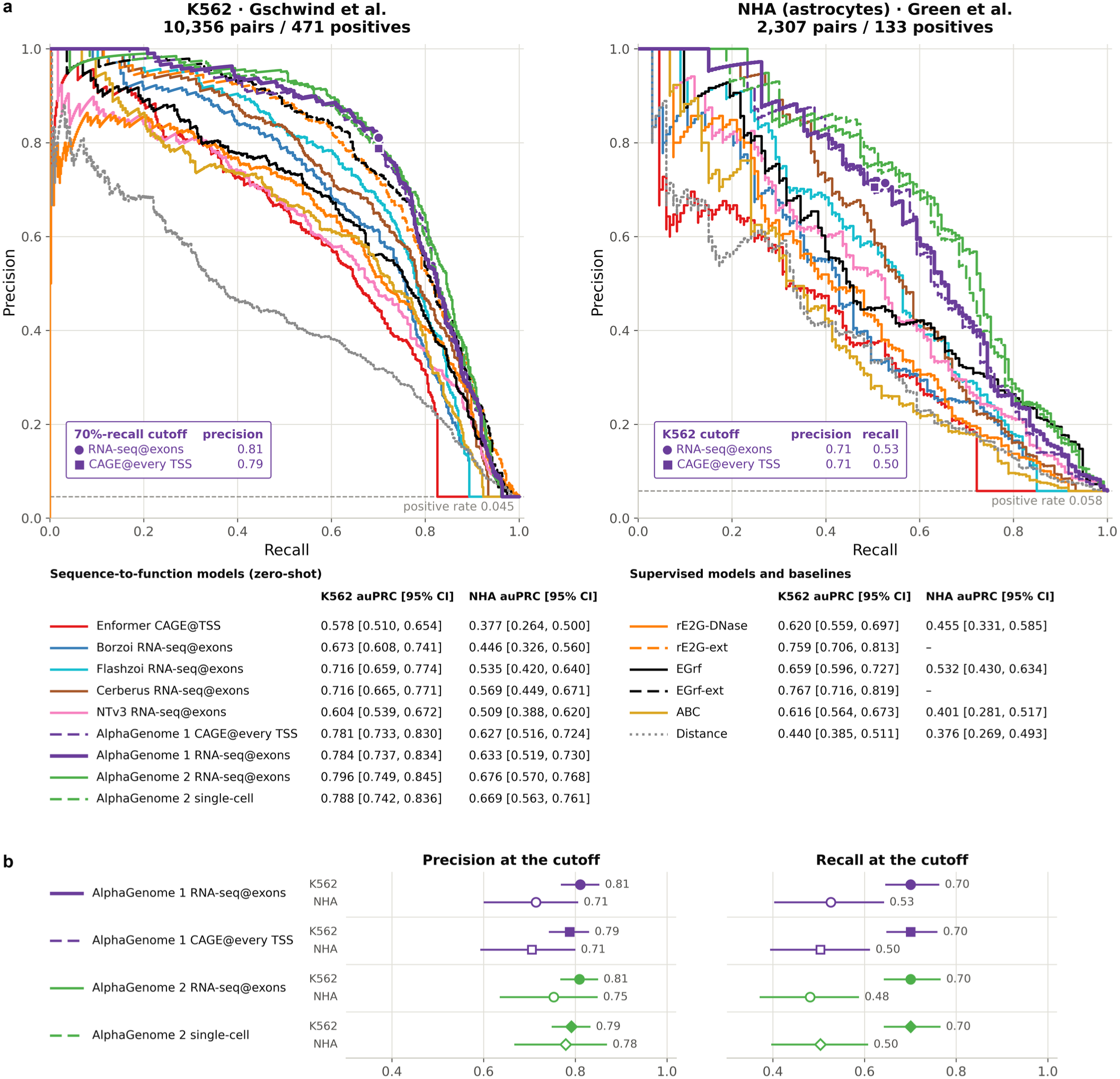
Precision–recall curves of selected methods, and precisions and recalls at the 70%-recall cutoff in the K562 screen. **a**, Precision–recall curves in K562 (left) and NHA (right) for selected seq2func readouts of Fig. 2, the expert models, and distance (1/distance to the MANE Select TSS), with auPRC [95% CI] as in Fig. 2. Markers, AlphaGenome 1 at the score that recovers 70% of the K562 CRISPRi positives (one cutoff per readout; circle, RNA-seq@exons; square, CAGE@every TSS), applied unchanged in NHA; boxes, their precision (and recall in NHA). On the released map’s own scores, the RNA-seq cutoff has a precision of 0.80 in K562 and 0.69 at a recall of 0.53 in astrocytes (Methods). Dashed lines, prevalence. **b**, Precision (left) and recall (right) at the same cutoff rule for the AlphaGenome 1 and AlphaGenome 2 readouts (shapes as in **a**; diamond, single-cell), in K562 (filled; recall 0.70 by construction) and NHA (open). Error bars, 95% confidence intervals (CI; connected-component bootstrap with the cutoff fixed, 10,000 replicates). No AlphaGenome 1 readout differed significantly from an AlphaGenome 2 readout in precision or recall (paired bootstrap; Benjamini–Hochberg within each screen, q ≥ 0.25; Supplementary Table 2). Pairs a method cannot score take a score of 0; ENCODE-rE2G and EGrf predictions are out-of-fold (held-out chromosomes).

**Supplementary Fig. 16.**
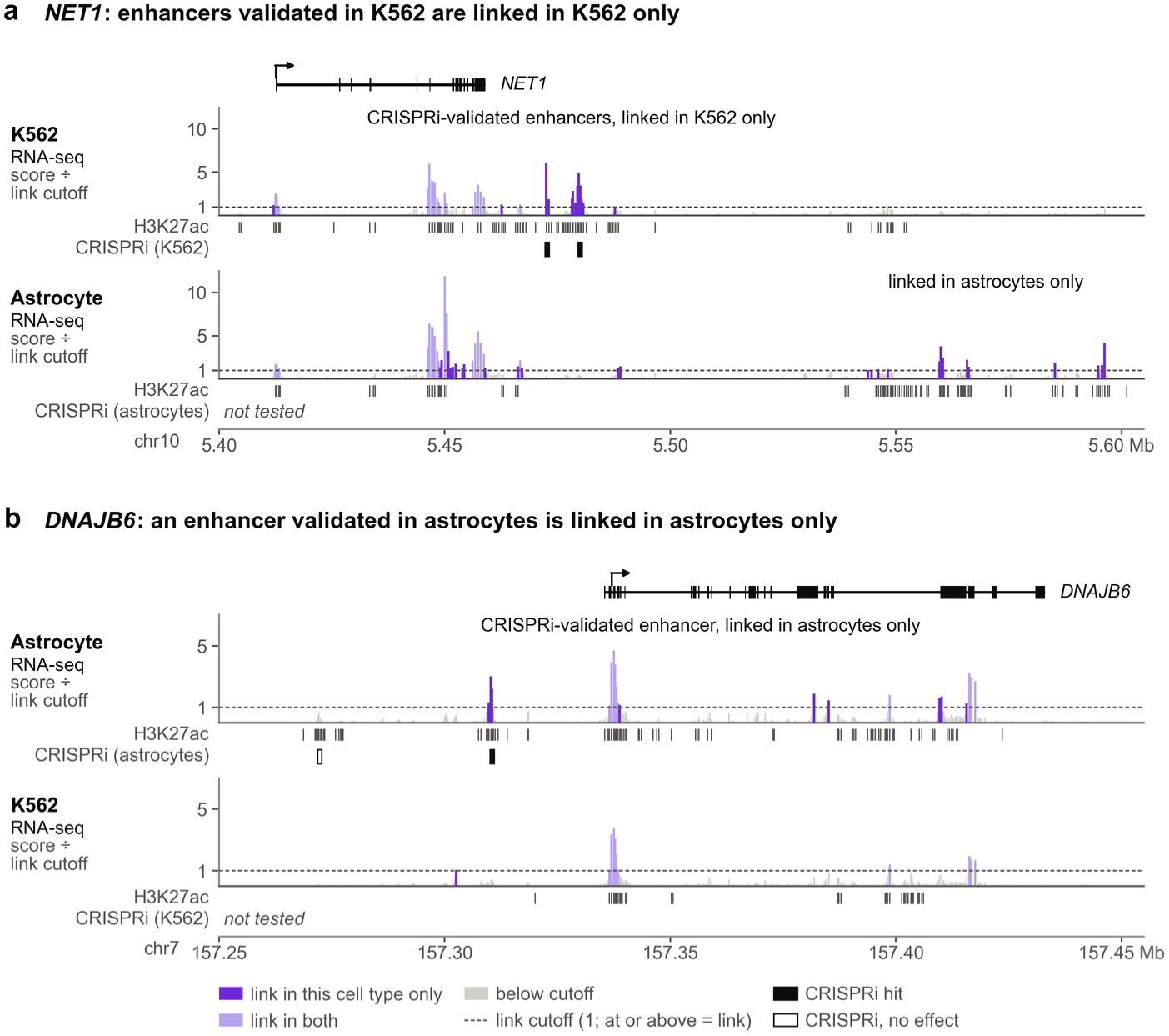
CRISPRi-validated enhancers pass the enhancer–gene link cutoff only in the screen’s cell type. E2GMap scores of registry cCREs in the default RNA-seq track of K562 and of astrocytes, divided by the RNA-seq link cutoff (the score at 70% recall of K562 CRISPRi positives; Methods). cCREs at or above 1 are links: dark, this cell type only; light, both; gray, below the cutoff. Filled boxes, CRISPRi hits; open box, tested element without effect. H3K27ac, ENCODE peaks. **a**, *NET1* (measured expression 120 TPM in K562 and 67 in astrocytes; ENCODE RNA-seq, as TPM among the map’s protein-coding genes), tested in K562 [9]: the four cCREs of its two hits score 1.9–6.1 times the cutoff in K562 and 0.29–0.52 times in astrocytes. **b**, *DNAJB6* (154 TPM in K562 and 133 in astrocytes), tested in astrocytes [10]: the cCRE of its hit scores 3.0 times the cutoff in astrocytes and 0.44 times in K562; the element without effect, 0.67 times in astrocytes.

## Notes

https://github.com/cthe4801/e2gscore

https://e2gscore.com

### Competing Interest Statement

The authors have declared no competing interest.

## References

1. Maurano, M. T. et al. Systematic localization of common disease-associated variation in regulatory DNA. Science 337, 1190–1195 (2012). 10.1126/science.1222794

2. Roadmap Epigenomics Consortium et al. Integrative analysis of 111 reference human epigenomes. Nature 518, 317–330 (2015). 10.1038/nature14248

3. Schoenfelder, S. & Fraser, P. Long-range enhancer–promoter contacts in gene expression control. Nat. Rev. Genet. 20, 437–455 (2019). 10.1038/s41576-019-0128-0

4. Gazal, S. et al. Combining SNP-to-gene linking strategies to identify disease genes and assess disease omnigenicity. Nat. Genet. 54, 827–836 (2022). 10.1038/s41588-022-01087-y

5. Fulco, C. P. et al. Activity-by-contact model of enhancer–promoter regulation from thousands of CRISPR perturbations. Nat. Genet. 51, 1664–1669 (2019). 10.1038/s41588-019-0538-0

6. Gasperini, M. et al. A genome-wide framework for mapping gene regulation via cellular genetic screens. Cell 176, 377–390.e19 (2019). 10.1016/j.cell.2018.11.029

7. Schraivogel, D. et al. Targeted Perturb-seq enables genome-scale genetic screens in single cells. Nat. Methods 17, 629–635 (2020). 10.1038/s41592-020-0837-5

8. Nasser, J. et al. Genome-wide enhancer maps link risk variants to disease genes. Nature 593, 238–243 (2021). 10.1038/s41586-021-03446-x

9. Gschwind, A. R. et al. An encyclopedia of human enhancer–gene regulatory interactions. Nature 657, 179–190 (2026). 10.1038/s41586-026-10781-4

10. Green, N. F. O. et al. CRISPRi screening in cultured human astrocytes uncovers distal enhancers controlling genes dysregulated in Alzheimer’s disease. Nat. Neurosci. 29, 703–716 (2026). 10.1038/s41593-025-02154-3

11. Ray, J., et al. A systematic survey of distal element-gene regulatory interactions with Direct-Capture Targeted Perturb-seq. Preprint at bioRxiv 10.1101/2025.09.16.676677 (2026).

12. Zhou, J. & Troyanskaya, O. G. Predicting effects of noncoding variants with deep learning–based sequence model. Nat. Methods 12, 931–934 (2015). 10.1038/nmeth.3547

13. Kelley, D. R. et al. Sequential regulatory activity prediction across chromosomes with convolutional neural networks. Genome Res. 28, 739–750 (2018). 10.1101/gr.227819.117

14. Zhou, J. et al. Deep learning sequence-based *ab initio* prediction of variant effects on expression and disease risk. Nat. Genet. 50, 1171–1179 (2018). 10.1038/s41588-018-0160-6

15. Sasse, A., Chikina, M. & Mostafavi, S. Unlocking gene regulation with sequence-to-function models. Nat. Methods 21, 1374–1377 (2024). 10.1038/s41592-024-02331-5

16. Zou, J. et al. A primer on deep learning in genomics. Nat. Genet. 51, 12–18 (2019). 10.1038/s41588-018-0295-5

17. Eraslan, G., Avsec, Ž., Gagneur, J. & Theis, F. J. Deep learning: new computational modelling techniques for genomics. Nat. Rev. Genet. 20, 389–403 (2019). 10.1038/s41576-019-0122-6

18. Consens, M. E., et al. Transformers and genome language models. Nat. Mach. Intell. 7, 346–362 (2025). 10.1038/s42256-025-01007-9

19. Novakovsky, G., Dexter, N., Libbrecht, M. W., Wasserman, W. W. & Mostafavi, S. Obtaining genetics insights from deep learning via explainable artificial intelligence. Nat. Rev. Genet. 24, 125–137 (2023). 10.1038/s41576-022-00532-2

20. Simonyan, K., Vedaldi, A. & Zisserman, A. Deep inside convolutional networks: visualising image classification models and saliency maps. Preprint at 10.48550/arXiv.1312.6034 (2014).

21. Toneyan, S. & Koo, P. K. Interpreting *cis*-regulatory interactions from large-scale deep neural networks. Nat. Genet. 56, 2517–2527 (2024). 10.1038/s41588-024-01923-3

22. Avsec, Ž., et al. Effective gene expression prediction from sequence by integrating long-range interactions. Nat. Methods 18, 1196–1203 (2021). 10.1038/s41592-021-01252-x

23. Linder, J., Srivastava, D., Yuan, H., Agarwal, V. & Kelley, D. R. Predicting RNA-seq coverage from DNA sequence as a unifying model of gene regulation. Nat. Genet. 57, 949–961 (2025). 10.1038/s41588-024-02053-6

24. Avsec, Ž., et al. Advancing regulatory variant effect prediction with AlphaGenome. Nature 649, 1206–1218 (2026). 10.1038/s41586-025-10014-0

25. Gayoso, A. et al. AlphaGenome 2 predicts regulatory activity and variant effects across cell states. Preprint at bioRxiv 10.64898/2026.11.11.765857 (2026).

26. Weber, L. M. et al. Essential guidelines for computational method benchmarking. Genome Biol. 20, 125 (2019). 10.1186/s13059-019-1738-8

27. Dehghani, M., et al. The benchmark lottery. Preprint at 10.48550/arXiv.2107.07002 (2021).

28. Karollus, A., Mauermeier, T. & Gagneur, J. Current sequence-based models capture gene expression determinants in promoters but mostly ignore distal enhancers. Genome Biol. 24, 56 (2023). 10.1186/s13059-023-02899-9

29. Huang, C. et al. Personal transcriptome variation is poorly explained by current genomic deep learning models. Nat. Genet. 55, 2056–2059 (2023). 10.1038/s41588-023-01574-w

30. Sasse, A. et al. Benchmarking of deep neural networks for predicting personal gene expression from DNA sequence highlights shortcomings. Nat. Genet. 55, 2060–2064 (2023). 10.1038/s41588-023-01524-6

31. Murphy, A. & Koo, P. K. Benchmarking seq2func models on distal enhancer effects with CRISPRi screens. Genomics x AI Blog 10.5281/zenodo.24198935 (2026).

32. Cheng, W. et al. DNALONGBENCH: a benchmark suite for long-range DNA prediction tasks. Nat. Commun. 16, 10108 (2025). 10.1038/s41467-025-65077-4

33. Huang, Z., Huang, R. & Han, J. Response representation shapes AlphaGenome enhancer-gene prediction in human astrocytes. Preprint at bioRxiv 10.64898/2026.08.14.744988 (2026).

34. Whalen, S., Truty, R. M. & Pollard, K. S. Enhancer–promoter interactions are encoded by complex genomic signatures on looping chromatin. Nat. Genet. 48, 488–496 (2016). 10.1038/ng.3539

35. Cao, Q. et al. Reconstruction of enhancer–target networks in 935 samples of human primary cells, tissues and cell lines. Nat. Genet. 49, 1428–1436 (2017). 10.1038/ng.3950

36. Xi, W. & Beer, M. A. Local epigenomic state cannot discriminate interacting and non-interacting enhancer–promoter pairs with high accuracy. PLoS Comput. Biol. 14, e1006625 (2018). 10.1371/journal.pcbi.1006625

37. Cao, F. & Fullwood, M. J. Inflated performance measures in enhancer–promoter interaction-prediction methods. Nat. Genet. 51, 1196–1198 (2019). 10.1038/s41588-019-0434-7

38. Whalen, S., Schreiber, J., Noble, W. S. & Pollard, K. S. Navigating the pitfalls of applying machine learning in genomics. Nat. Rev. Genet. 23, 169–181 (2022). 10.1038/s41576-021-00434-9

39. Steegen, S., Tuerlinckx, F., Gelman, A. & Vanpaemel, W. Increasing transparency through a multiverse analysis. Perspect. Psychol. Sci. 11, 702–712 (2016). 10.1177/1745691616658637

40. Hingerl, J. C., Karollus, A. & Gagneur, J. Flashzoi: an enhanced Borzoi for accelerated genomic analysis. Bioinformatics 41, btaf467 (2025). 10.1093/bioinformatics/btaf467

41. Boshar, S., et al. A multispecies foundation model for genome function prediction, gene annotation and sequence design. Preprint at bioRxiv 10.64898/2025.12.22.695963 (2026).

42. Kelley, D. R., Yuan, H., Huang, X. & Linder, J. Cerberus: bidirectional state space blocks improve accuracy and efficiency of regulatory sequence models. Preprint at bioRxiv 10.64898/2026.10.02.756377 (2026).

43. Morales, J. et al. A joint NCBI and EMBL-EBI transcript set for clinical genomics and research. Nature 604, 310–315 (2022). 10.1038/s41586-022-04558-8

44. Benjamini, Y. & Hochberg, Y. Controlling the false discovery rate: a practical and powerful approach to multiple testing. J. R. Stat. Soc. B 57, 289–300 (1995). 10.1111/j.2517-6161.1995.tb02031.x

45. Mudge, J. M., et al. GENCODE 2025: reference gene annotation for human and mouse. Nucleic Acids Res. 53, D966–D975 (2025). 10.1093/nar/gkae1078

46. Moore, J. E. et al. An expanded registry of candidate *cis*-regulatory elements. Nature 10.1038/s41586-025-09909-9 (2026).

47. Sheth, M. U. et al. Mapping enhancer–gene regulatory interactions from single-cell data. Nat. Genet. 58, 1941–1952 (2026). 10.1038/s41588-026-02695-8

48. Nair, S., Shrikumar, A., Schreiber, J. & Kundaje, A. fastISM: performant *in silico* saturation mutagenesis for convolutional neural networks. Bioinformatics 38, 2397–2403 (2022). 10.1093/bioinformatics/btac135

49. Bredikhin, D., et al. Porting AlphaGenome to PyTorch. Genomics x AI Blog 10.5281/zenodo.20272906 (2026).

50. Majdandzic, A., Rajesh, C. & Koo, P. K. Correcting gradient-based interpretations of deep neural networks for genomics. Genome Biol. 24, 109 (2023). 10.1186/s13059-023-02956-3

51. Altschul, S. F. & Erickson, B. W. Significance of nucleotide sequence alignments: a method for random sequence permutation that preserves dinucleotide and codon usage. Mol. Biol. Evol. 2, 526–538 (1985). 10.1093/oxfordjournals.molbev.a040370

52. Deming, W. E. & Stephan, F. F. On a least squares adjustment of a sampled frequency table when the expected marginal totals are known. Ann. Math. Stat. 11, 427–444 (1940). 10.1214/aoms/1177731829

53. The ENCODE Project Consortium, et al. Expanded encyclopaedias of DNA elements in the human and mouse genomes. Nature 583, 699–710 (2020). 10.1038/s41586-020-2493-4

54. The GTEx Consortium. The GTEx Consortium atlas of genetic regulatory effects across human tissues. Science 369, 1318–1330 (2020). 10.1126/science.aaz1776

55. The FANTOM Consortium and the RIKEN PMI and CLST (DGT). A promoter-level mammalian expression atlas. Nature 507, 462–470 (2014). 10.1038/nature13182

